# Evolutionary branching in distorted trait spaces

**DOI:** 10.1101/794966

**Authors:** Hiroshi C. Ito, Akira Sasaki

**Affiliations:** Department of Evolutionary Studies of Biosystems, The Graduate University for Advanced Studies, SOKENDAI, Hayama, Kanagawa 240-0193, Japan; Evolution and Ecology Program, International Institute for Applied Systems Analysis, Laxenburg, Austria

## Abstract

Biological communities are thought to have been evolving in trait spaces that are not only multi-dimensional, but also distorted in a sense that mutational covariance matrices among traits depend on the parental phenotypes of mutants. Such a distortion may affect diversifying evolution as well as directional evolution. In adaptive dynamics theory, diversifying evolution through ecological interaction is called evolutionary branching. This study analytically develops conditions for evolutionary branching in distorted trait spaces of arbitrary dimensions, by a local nonlinear coordinate transformation so that the mutational covariance matrix becomes locally constant in the neighborhood of a focal point. The developed evolutionary branching conditions can be affected by the distortion when mutational step sizes have significant magnitude difference among directions, i.e., the eigenvalues of the mutational covariance matrix have significant magnitude difference.

## 1 Introduction

Biological communities are thought to have been evolving in multi-dimensional trait spaces (Doebeli and Ispolatov, 2010, 2017), and which are distorted in a sense that mutatability in each direction (i.e., the mutational covariance matrix) varies depending on the parental phenotype of the mutant. Such a distortion may affect evolutionary dynamics and outcomes, including directional evolution and diversifying evolution.

Directional evolution in distorted trait spaces can be described with an ordinary differential equation for the resident trait, derived under assumption of the rare and small mutation limit in adaptive dynamics theory (Dieckmann and Law, 1996), or that for the mean trait under some assumption on variances and on higher moments of the trait in quantitative genetics (Lande, 1979). In both frameworks, directional evolution is shown to be proportional to the fitness gradient (or selection gradient) multiplied by the mutational covariance matrix (or additive genetic covariance matrix). In distorted trait spaces, the covariance matrix varies depending on the parental phenotypes of mutants, which can change the speed and/or direction of directional evolution (explained in Section 2.1).

Diversifying evolution, which is a fundamental source of biodiversity, is described as evolutionary branching in adaptive dynamics theory (Metz et al., 1996; Geritz et al., 1997). Evolutionary branching explains sympatric and parapatric speciation as continuous adaptive evolution through ecological interaction (Dieckmann and Doebeli, 1999; Doebeli and Dieckmann, 2003; Dieckmann et al., 2004; Doebeli 2011). If a space composed of evolutionary traits has an evolutionary branching point, the point attracts a monomorphic population through directional selection, and then favors its diversification through disruptive selection (Geritz et al., 1997). Moreover, a trait space may have not only evolutionary branching points but also evolutionary branching lines (Ito and Dieckmann, 2012, 2014), if the trait space has a significant mutatability difference among directions so that mutation in some direction is significantly difficult compared to the other directions. Analogously to evolutionary branching points, an evolutionary branching line attracts a monomorphic population and then favors their evolutionary branching through disruptive selection (Ito and Dieckmann, 2014).

So far, in multi-dimensional trait spaces, existence conditions for evolutionary branching points and lines (for two-dimensional trait spaces), or candidates for them (for the higher-dimensions, explained in Appendix D), have been developed under assumption of constant mutational covariance matrices, i.e., no distortion. Thus, a next step is to reveal these conditions in distorted trait spaces.

In this paper, we first in the literature analyze the evolutionary branching in distorted trait spaces. By means of a local coordinate normalization to make the distortion vanish locally, we formally develop the conditions for evolutionary branching points and lines for two-dimensional distorted trait spaces. Although the analogous conditions are obtained in distorted trait spaces of arbitrarily higher dimensions (Appendix D), for simplicity, we restrict our explanation to two-dimensional trait spaces in the main text. For convenience, we refer to the conditions for evolutionary branching points and lines as the branching point conditions and branching line conditions, respectively.

To show with a minimum complexity how the distortion of a trait space affects evolutionary branching, Section 2 considers a simply distorted trait space and derives the branching point conditions and branching line conditions. Section 3 derives analogous results in an arbitrarily distorted trait space. Section 4 is devoted to an example to show how this theory can be applied. Section 5 discusses the obtained results in connection with relevant studies.

## 2 Evolutionary branching in a simply distorted trait space

Throughout the paper, we use italic for denoting scalars, bold small letters for column vectors, and bold capital for matrices. We consider a two-dimensional trait space **s** = (*x*, *y*)^T^ and a monomorphic population with a resident phenotype **s** = (*x*, *y*)^T^, where T denotes transpose. From resident **s**, a mutant **s**′ = (*x*′, *y*′)^T^ emerges with a mutation rate *μ* per birth event. The point **s**′ a mutant resides in the trait space follows a probability density distribution *m*(**s**′, **s**), referred to as a “mutation distribution.” The shape of *m*(**s**′, **s**) changes depending on **s**.

### 2.1 Adaptive dynamics theory

To analyze adaptive evolution in the trait space **s** = (*x*, *y*)^T^, we use one of adaptive dynamics theories, which is originated from Metz et al. (1996). This theory typically assumes clonal reproduction, sufficiently rare mutation, and sufficiently large population size, so that a population is monomorphic and is almost at an equilibrium density whenever a mutant emerges. In this setting, whether a mutant can invade the resident is determined by its initial per capita growth rate, called the invasion fitness, *f*(**s**′, **s**), which is a function of mutant **s**′ and resident **s**. The invasion fitness *f*(**s**′, **s**) can be translated into a fitness landscape along mutant trait **s**′. The landscape can vary depending on the resident trait **s**. The mutant can invade the resident only when *f*(**s**′, **s**) is positive, resulting in substitution of the resident in many cases. Repetition of such a substitution forms directional evolution toward a higher fitness, as long as the dominant component of the fitness landscape around **s** is the fitness gradient (corresponding to directional selection) rather than the fitness curvature (corresponding to diversifying or purifying selection). When the fitness gradient becomes small so that the second-order fitness component is not negligible, a mutant may coexist with its resident, which may bring about evolutionary diversification into two distinct morphs, called evolutionary branching (Metz et al., 1996; Geritz et al., 1997; Geritz et al., 1998). Such an evolutionary transition of residents induced by repeated mutant invasions, including directional evolution and evolutionary branching, is called a trait substitution sequence (Metz et al., 1996).

The expected evolutionary shift of resident phenotype through directional evolution is approximated as

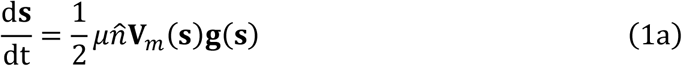

(Dieckmann and Law, 1996), where *μ* is the mutation rate, 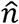 is the equilibrium population density of the resident **s**, **V**_*m*_(**s**) is the covariance matrix of the mutation distribution *m*(**s**′, **s**), and

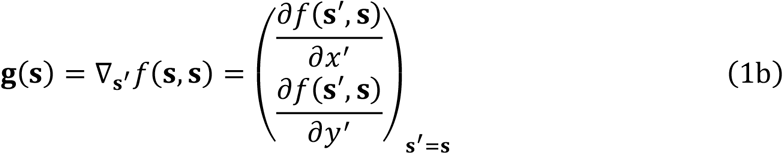

is the fitness gradient vector evaluated at the resident trait **s**. Eqs. (1) are applicable even when the mutation distribution varies over **s** and so does **V**_*m*_(**s**). In this case, such a dependency affects not only the speed of directional evolution but also its direction (Fig.1).

**Figure 1.**
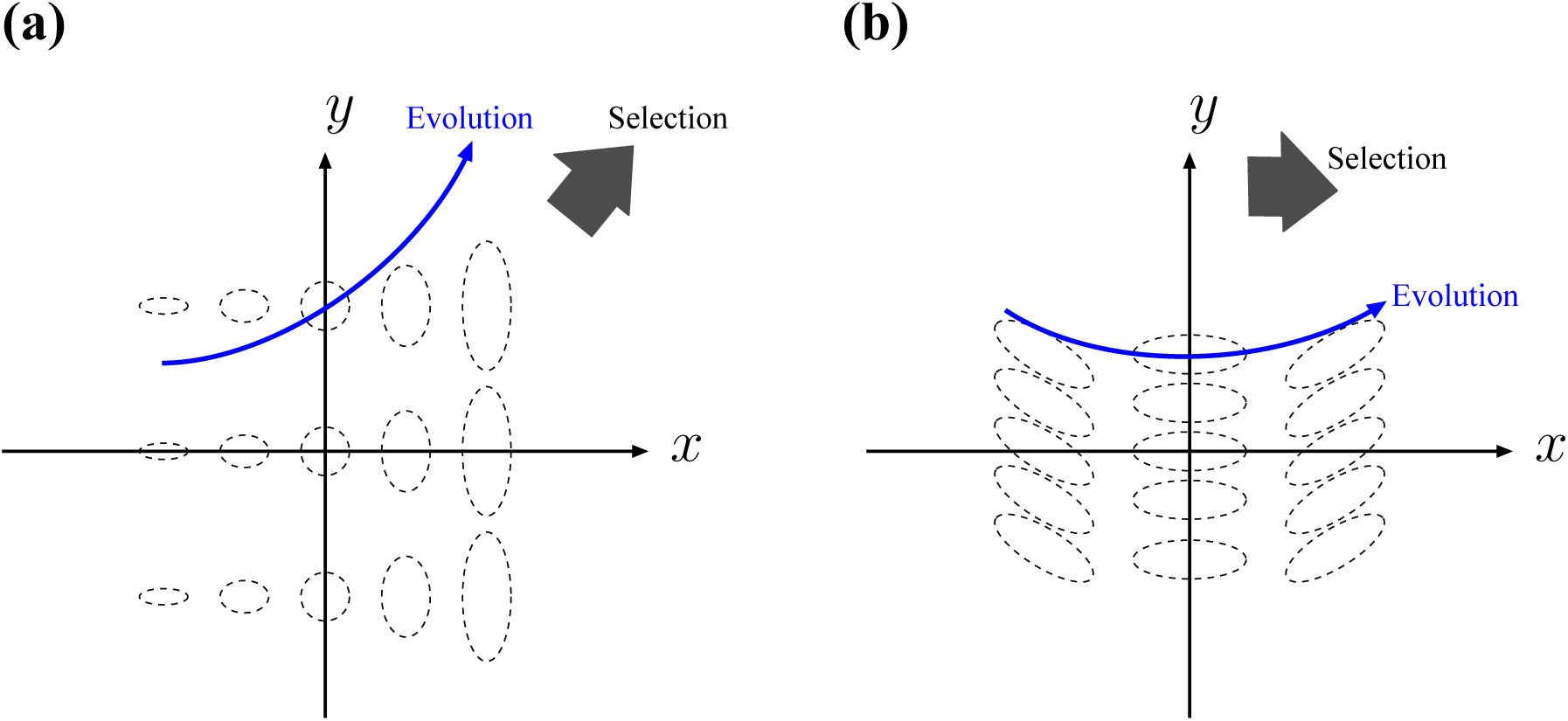
Illustrated directional evolution affected by distortion of trait space. In panels (a) and (b), the covariance matrices of mutation distributions, indicated with black dotted ellipses, vary depending on resident phenotypes (i.e., trait spaces are distorted). In both cases, directionally evolving populations described with Eqs. (1) are expected to change their directions (blue curved arrows) as well as speeds, even under constant selection gradients (dark gray arrows).

### 2.2 Assumption for mutation

As for evolutionary branching in two-dimensional trait spaces, branching point conditions (Geritz et al., 2016) and branching line conditions (Ito and Dieckmann, 2012, 2014) are applicable only for non-distorted trait spaces (i.e., the mutation distribution does not depend on the resident phenotype). To apply those branching conditions for distorted trait spaces, we assume there exists a nonlinear transformation of the coordinate system **s** = (*x*, *y*)^T^ into a new coordinate system 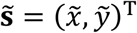 in which the mutation distribution can be approximated with a bivariate Gaussian distribution that is constant at least locally around a focal point **s**_0_. We refer to the coordinates **s** = (*x*, *y*)^T^ and 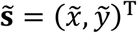 as the “original coordinates” and “geodesic coordinates”, respectively (the meaning of “geodesic” is explained in Section 3.1). However, if we approximate the mutation distribution with a Gaussian distribution in the original coordinates **s** (with its covariance matrix given by **V**_*m*_(**s**)), the nonlinear coordinate transformation can cause the mutation distribution in the geodesic coordinates 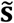 to deviate from the Gaussian. Such a deviation is not negligible when the mutation distribution has a significantly narrow width in one direction compared to the other (Fig. 2a). In this case, the branching line conditions are not applicable even in the geodesic coordinates 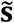. To avoid this difficulty, in this paper we assume the Gaussian approximation not before but after the coordinate transformation (Fig. 2b). This choice has an advantage that we can express mutants restricted to constraint curves completely or incompletely (left panel in Fig. 2b).

**Figure 2.**
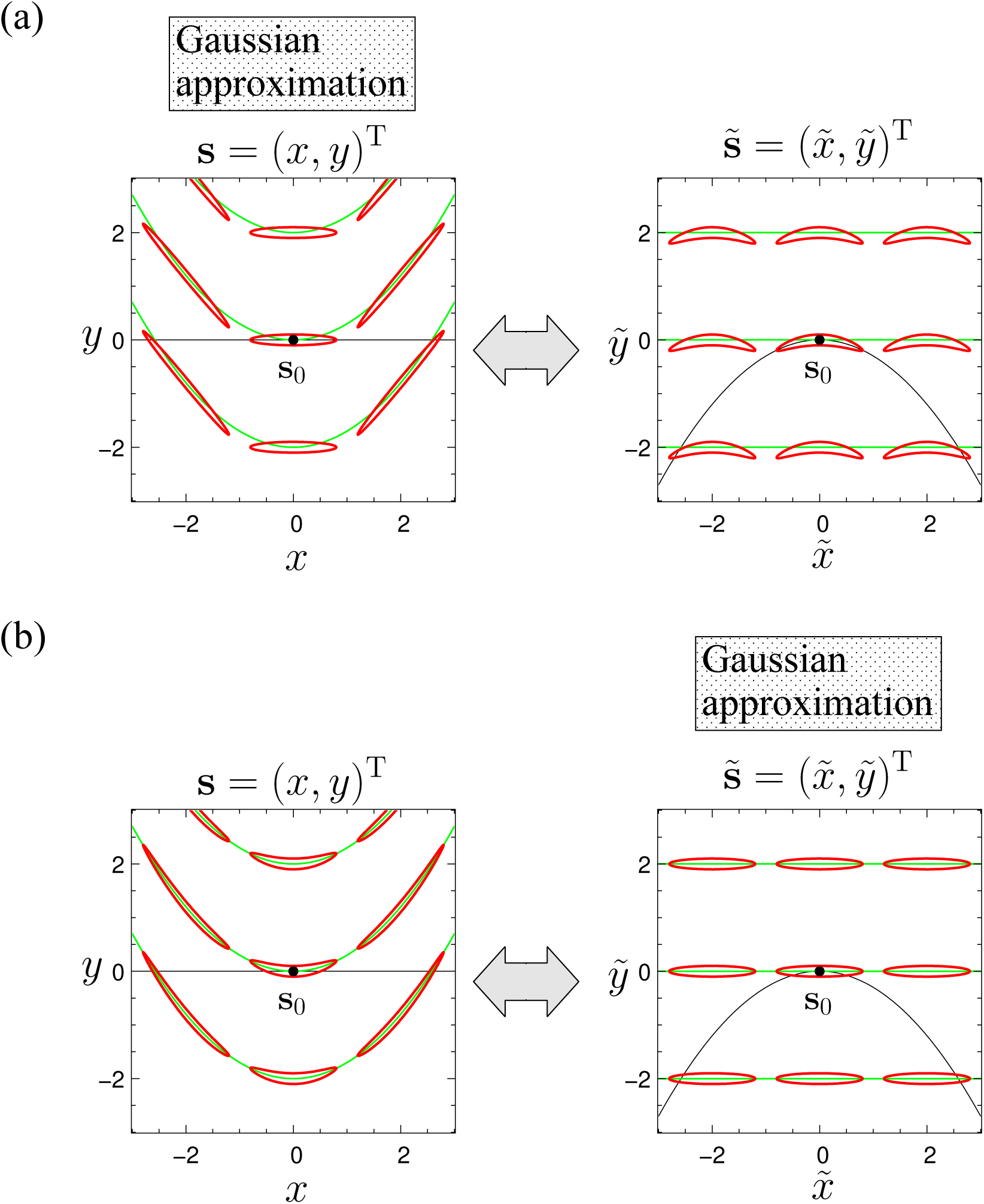
Gaussian approximation of mutation distributions before or after nonlinear coordinate transformation. Red closed curves are contours of mutation distributions defined by Eq. (14b), referred to as mutation contours. Green curves indicate constraint curves formed under *σ*_*y*_ = 0. The coordinate transformation is defined by Eqs. (2). Parameters: *σ*_*x*_ = 0.8, *σ*_*y*_ = 0.1, and *ρ* = 0.6.

To show with a minimal complexity how distortion of a trait space affects evolutionary branching conditions, we assume that the nonlinear transformation from the original coordinates **s** = (*x*, *y*)^T^ (around the focal point **s**_0_ = (*x*_0_, *y*_0_)^T^) into the geodesic coordinates 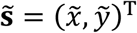 is given by

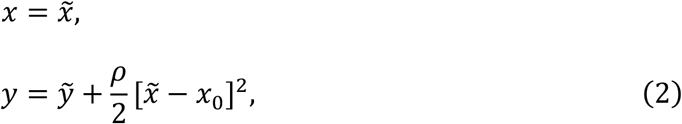

with a single parameter *ρ* for controlling the degree of distortion (Fig. 2b). In addition, we assume that the mutation distribution in the geodesic coordinates 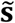 is a constant bivariate Gaussian distribution,

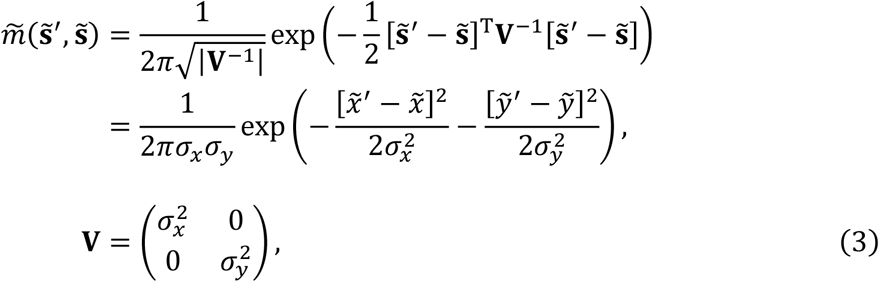

as illustrated in the right side of Fig. 2b. Then, expressing 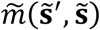 in the original coordinates **s** gives the mutation distribution *m*(**s**′, **s**) that varies depending on **s**, as illustrated in the left side of Fig. 2b (see Appendix A.1 for the derivation of *m*(**s**′, **s**)). The *σ*_*x*_ and *σ*_*y*_ describe the standard deviations of mutation along the 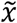- and 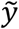-directions, respectively. When *σ*_*y*_ is very small, mutants deriving from an ancestral resident 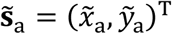 are almost restricted to a line 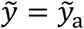 (i.e., 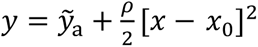), but can deviate slightly from it (Fig. 2b).

The local distortion defined by Eqs. (2) and (3) is a special case that is much simpler than a general expression for the local distortion defined by Eqs. (11) in the next section. However, the branching point conditions and branching line conditions derived in this simple case are essentially the same with those in the general case (Section 3.3–3.4). In this sense, the special case analyzed here has a certain generality.

By substituting Eqs. (2) into the invasion fitness function *f*(**s**′, **s**) in the original coordinates **s**, we obtain the invasion fitness function in the geodesic coordinates 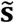, referred to as the “geodesic invasion fitness”,

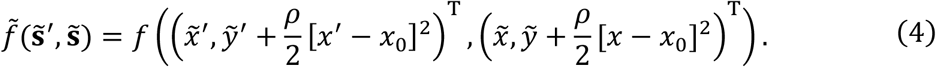

Note that the constant Gaussian mutation distribution in the geodesic coordinates 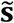 allows application of the branching point conditions and branching line conditions. The contribution of *ρ* on those conditions shows how distortion of the trait space affects them.

### 2.3 Quadratic approximation of invasion fitness functions

Both of the branching point conditions and branching line conditions depend only on the first and second derivatives of invasion fitness functions with respect to mutant and resident phenotypes. Thus, to facilitate analysis, we apply quadratic approximation to the original and geodesic invasion fitness functions, *f*(**s**′, **s**) and 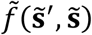, without loss of generality. Since the resident phenotype is at population dynamical equilibrium, *f*(**s**, **s**) = 0 must hold for any **s**. Then, following Ito and Dieckmann (2014), we expand *f*(**s**′, **s**) around the focal point **s**_0_ in the form of

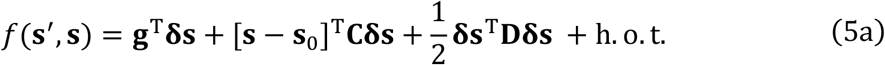

with **δs** = **s**′ − **s**,

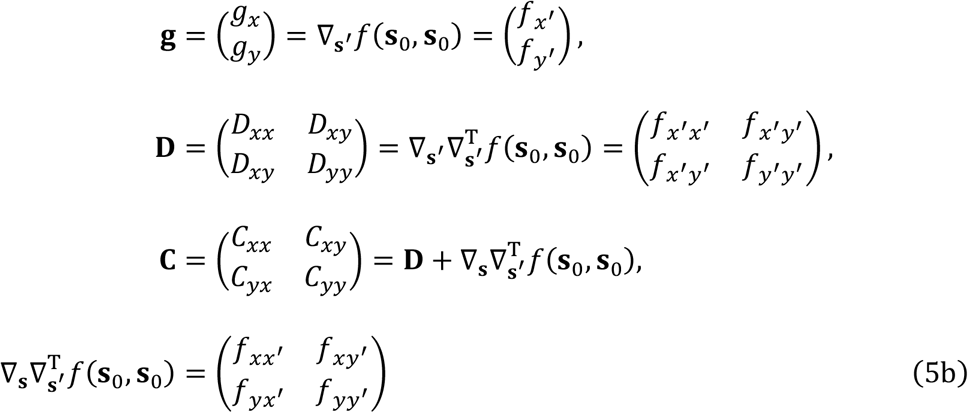

(Appendix B), where *f*_*α*_ = *∂f*(**s**′, **s**)/∂α for *α* = *x*′, *y*′, *x*, *y* and *f*_*αβ*_ = *∂*^2^*f*(**s**′, **s**)/ *∂α∂β* for *α, β* = *x*′, *y*′, *x*, *y* denote the first and second derivatives of *f*(**s**′, **s**), respectively, evaluated at **s**′ = **s** = **s**_0_. Note that *f*(**s**′, **s**) can be treated as a fitness landscape along **s**′, which varies depending on **s**. When g = 0, i.e., the point **s**_0_ is evolutionary singular (Geritz et al., 1997), the curvature of the fitness landscape along a vector v is given by **v**^T^**Dv**/|**v**|^2^. In other words, the signs of the two real eigenvalues of **D** determines whether the point **s**_0_ is a mountain top (locally evolutionarily stable (Maynard Smith and Price, 1973)), a basin bottom (evolutionarily unstable in all directions), or a saddle point (evolutionarily unstable in some directions). Even when **g** ≠ **0**, the sign of **v**^T^**Dv**/|**v**|^2^ tells whether the fitness landscape is locally convex or concave along v. In this sense, we refer to **D** as the “fitness curvature”.

For resident **s** deviated slightly from the focal point **s**_0_, the fitness gradient at **s** is approximately given by

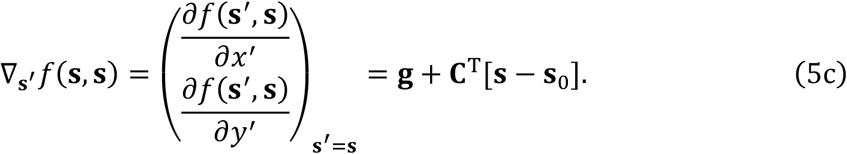

Thus, the matrix **C** describes the change rate of the fitness gradient when the resident deviates from **s**_0_. In this sense, we refer to **C** as the “fitness gradient variability.” When **g** = **0**, the Jacobian matrix **J** = **V**_*m*_(**s**_0_)**C**^T^ determines the local stability of **s**_0_ through directional evolution described by Eqs. (1) with Eq. (5c). Thus, if the all eigenvalues of **J** have negative real parts, then the point **s**_0_ is locally stable through directional evolution. Whenever the symmetric part of **C** is negative definite, the all eigenvalues of **J** have negative real parts as far as **V**_*m*_(**s**_0_) is non-singular (i.e., mutation is possible in all directions), in which case **s**_0_ is called a strongly convergence stable point (Leimar, 2009).

Substituting Eqs. (2) into Eqs. (5) gives the quadratic form for the geodesic invasion fitness function,

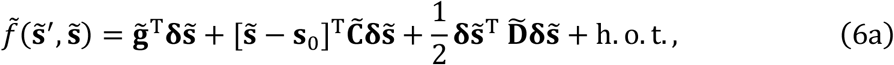

with 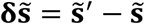,

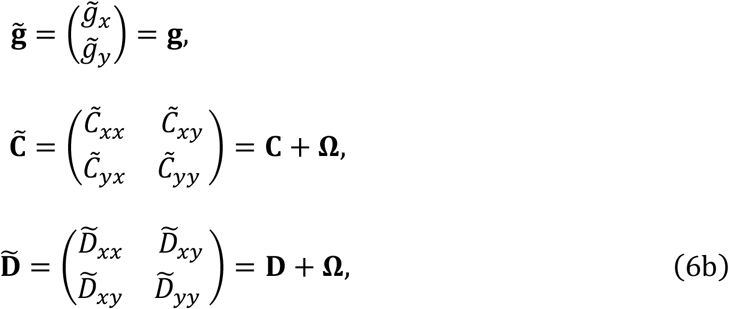

and

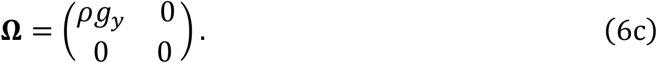

Since 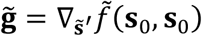, 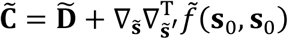 and 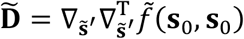 hold, they respectively describe the fitness gradient, fitness gradient variability, and fitness curvature at the focal point **s**_0_ in the geodesic coordinates 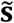. Note that **C** and **D** in the original coordinates **s** are respectively integrated with the “distortion effect” **Ω**, into 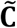 and 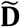 in the geodesic coordinates 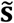. On the basis of the local coordinate normalization above, we derive the conditions for the focal point **s**_0_ being an evolutionary branching point (branching point conditions), and the conditions for existence of an evolutionary branching line containing **s**_0_ (branching line conditions), in the following subsections.

### 2.4 Conditions for evolutionary branching points

An evolutionary branching point attracts a monomorphic population in its neighborhood through directional evolution, and then favors its diversification into two morphs that directionally evolve in opposite directions (Metz et al., 1996; Geritz et al., 1997). Conditions for existence of evolutionary branching points, i.e., branching point conditions, have been derived originally in one-dimensional trait spaces (Geritz et al., 1997). For two-dimensional non-distorted trait spaces, the branching point conditions have been proved by approximating the latter diversification process with coupled Lande equations (Geritz et al. 2016). By expressing these two-dimensional branching point conditions in the geodesic coordinates 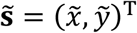, we derive the branching point conditions for the simply distorted trait space **s** = (*x*, *y*)^T^. Specifically, we obtain the following conditions for the focal point **s**_0_ being an evolutionary branching point.

i. **s**_0_ is evolutionarily singular, satisfying

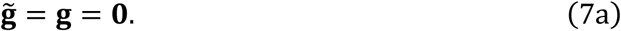
ii. **s**_0_ is strongly convergence stable, i.e., the symmetric part of

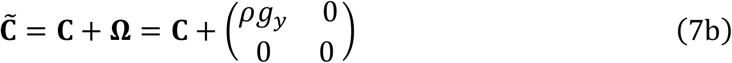

is negative definite.
iii. **s**_0_ is evolutionarily unstable, i.e., a symmetric matrix

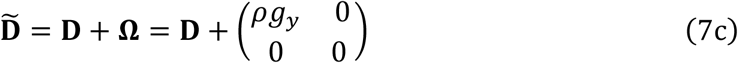

has at least one positive eigenvalue, in which case the fitness landscape is concave along at least one direction.

Since Eq. (7a) requires *g*_*x*_ = *g*_*y*_ = 0, we see 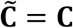 and 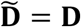. This means that the branching point conditions in the geodesic coordinates 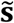 are equivalent to those in the original coordinates **s**. Thus, the simple distortion of the trait space, controlled by *ρ* in Eqs. 2, does not affect the branching point conditions.

### 2.5 Conditions for evolutionary branching lines

As long as *σ*_*y*_ has a comparable magnitude with *σ*_*x*_, evolutionary branching is expected only around the evolutionary branching points (Ito and Dieckmann, 2014). On the other hand, if *σ*_*y*_ is extremely smaller than *σ*_*x*_, the resulting slower evolutionary change in 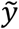 is negligible during the faster evolution in 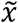, so that the evolutionary dynamics in the faster time scale can be described in a one-dimensional trait space 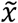 under fixed 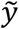. In this case, a point satisfying the one-dimensional conditions for evolutionary branching points (Geritz, et al. 1997) in 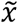 can induce evolutionary branching in 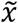. Even if *σ*_*y*_ is not extremely small, this type of evolutionary branching is likely to occur, as long as the disruptive selection along 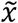, measured by 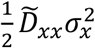, is sufficiently stronger than the directional selection along 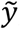, measured by 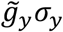 (Ito and Dieckmann, 2007, 2014). The conditions for this type of evolutionary branching are called the conditions for evolutionary branching lines or the branching line conditions, because points that satisfy those conditions often form lines in trait spaces, called evolutionary branching lines (Ito and Dieckmann, 2014).

To facilitate application of the branching line conditions, we simplify the original branching line conditions, following Ito and Dieckmann (2014) (see Appendix C.1-3 for details of the original branching line conditions and the simplification). Specifically, when *σ*_*y*_ is much smaller than *σ*_*x*_ so that 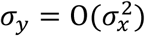 with *σ*_*x*_ ≪ 1 (i.e., *σ*_*y*_ has no larger magnitude than 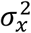), following Ito and Dieckmann (2014), we can further simplify Eq. (6a) into

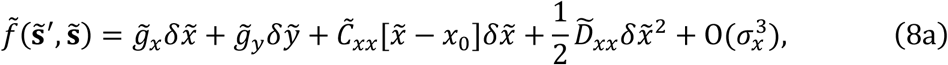

with

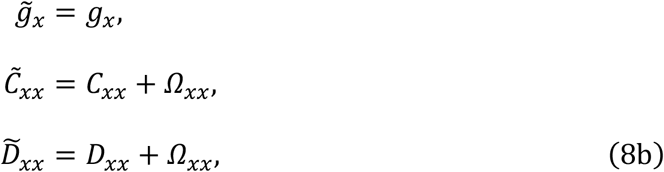

and

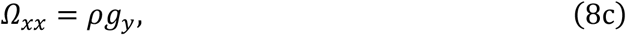

where terms with 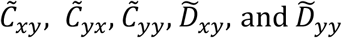, and 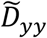 are subsumed in 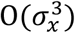. Note that this simplification is allowed even when *σ*_*y*_ is not much smaller than *σ*_*x*_, as long as magnitudes of 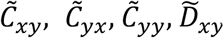, and 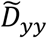 are all sufficiently small instead. According to Appendix B in Ito and Dieckmann (2014), Eqs. (8) hold when the sensitivity of the geodesic invasion fitness, 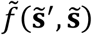, to single mutational changes of 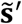 and 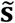 is significantly lower in 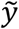 than in 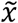, satisfying

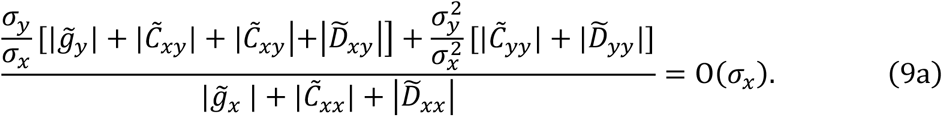

On this basis, the simplified branching line conditions are described as follows:

i. At **s**_0_ the sensitivity of 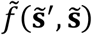 to single mutational changes of 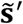 and 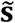 is significantly lower in 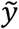 than in 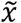, satisfying Eq. (9a).
ii. **s**_0_ is evolutionarily singular along 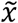, satisfying

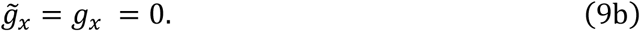
iii. **s**_0_ is convergence stable along 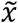, satisfying

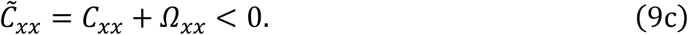
iv. **s**_0_ is sufficiently evolutionarily unstable (i.e., subject to sufficiently strong disruptive selection) along 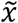, satisfying

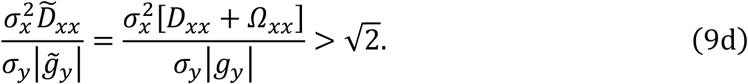

Note that condition (ii) above does not require *g*_*y*_ = 0, and thus *Ω*_*xx*_ = *ρg*_*y*_ may remain nonzero in Eqs. (9c) and (9d). Thus, differently from the branching point conditions, distortion of the trait space affects the branching line conditions through *Ω*_*xx*_ = *ρg*_*y*_, as long as the fitness gradient along the *y*-axis, *g*_*y*_, exists.

Existence of an evolutionary branching line ensures the occurrence of evolutionary branching of a monomorphic population located in its neighborhood, in the maximum likelihood invasion-event path, i.e., a trait substitution sequence composed of mutant-invasion events each of which has the maximum likelihood (Ito and Dieckmann 2014).

When *σ*_*y*_ = 0, the evolutionary trajectory starting from the focal point **s**_0_ in the geodesic coordinates 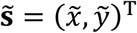 is completely restricted to the line 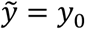, which forms a parabolic curve in the original coordinates **s** = (*x*, *y*)^T^,

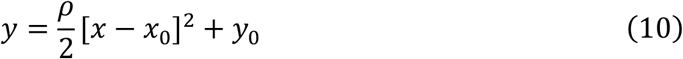

(green curves in Fig. 2b). In this case, condition (i) always holds and condition (iv) is simplified into 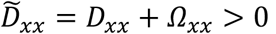, and thus conditions (ii-iv) become the one-dimensional branching point conditions (Geritz et al., 1997) in 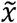 treated as a one-dimensional trait space. In the original coordinates **s** = (*x*, *y*)^T^, conditions (ii-iv) give the conditions for evolutionary branching point along a constraint curve locally approximated in the form of Eq. (10), and which are identical to the three conditions derived by Ito and Sasaki (2016) with an extended Lagrange multiplier method. Thus, the above conditions with *σ*_*y*_ > 0 extend the conditions by Ito and Sasaki (2016) for the case allowing slight mutational deviations from the constraint curves, i.e., when the constraints are incomplete.

Although this section focuses on one of the simplest configurations among possible local distortions for two-dimensional trait spaces, the obtained results are already useful in analyses of eco-evolutionary models defined on two-dimensional trait spaces with constraint curves deriving from various trade-offs (e.g., trade-offs between competitive ability and grazing susceptibility of primary producers (Branco et al., 2010), foraging gain and predation risk of consumers (Abrams, 2003), specialist and generalist of consumers (Egas et al., 2004), transmission and virulence of parasites (Kamo et al., 2006), competitive ability and attack rate (or longevity) of parasitoids (Bonsal et al., 2004), and fecundity and dispersal (Weigang and Kisdi, 2015)). Specifically, by an appropriate rotation around a focal point (Fig. 3a to 3b) and obtaining the geodesic coordinates (Fig. 3b to 3c), we can apply the branching line conditions, Eqs. (9), which tells likelihoods of evolutionary branching in those models when the constraint curves are incomplete as well as complete.

**Figure 3.**
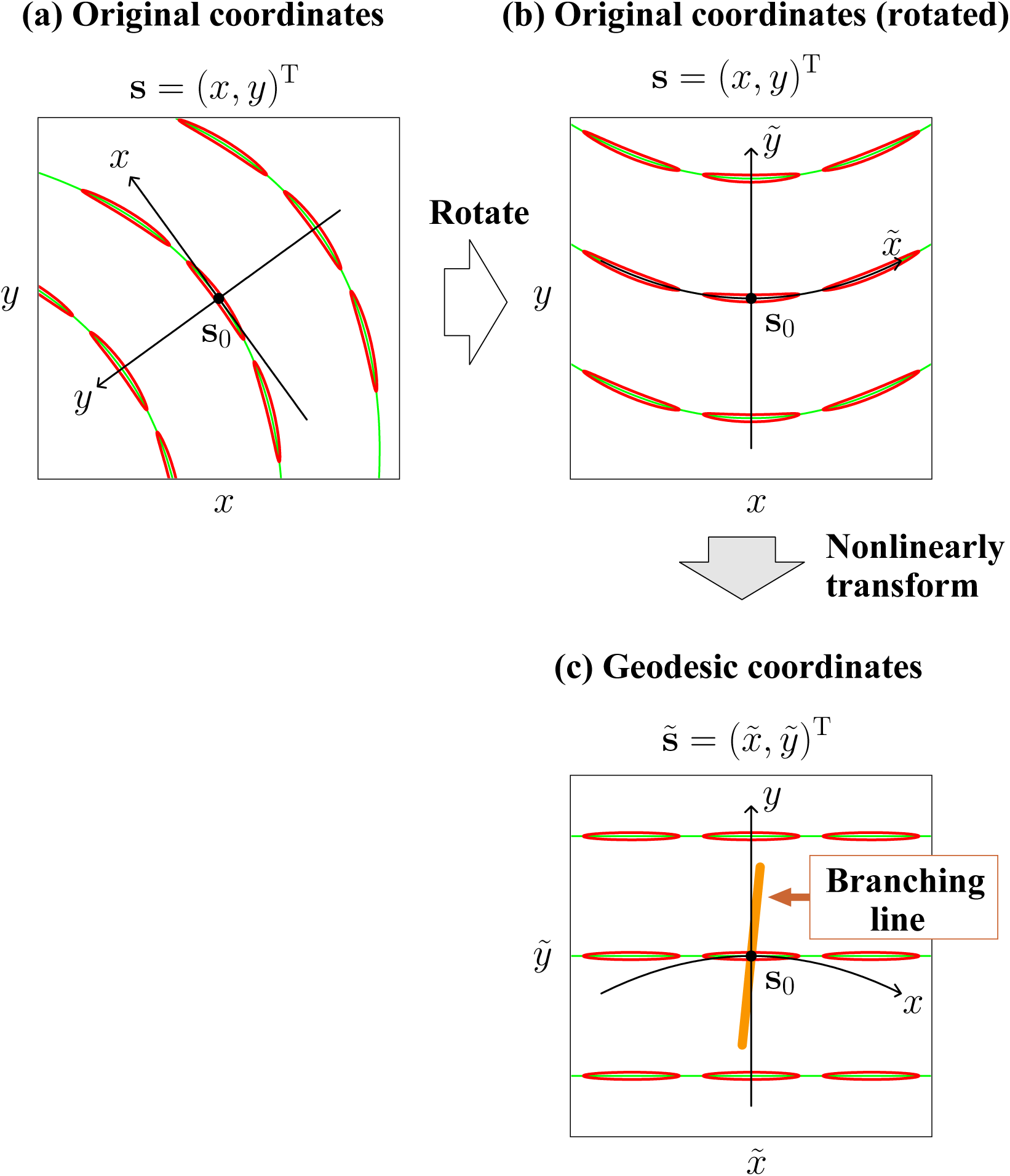
Illustrated application of branching line conditions derived for simply distorted trait spaces in Section 2.5. In eco-evolutionary models defined on two-dimensional trait spaces with constraint curves, we can analyze the likelihood of evolutionary branching not only when all mutants are completely restricted to the curves (complete constraints), but also when some mutants can slightly deviate from the curves (incomplete constraints). The branching line conditions can be applied in the geodesic coordinates, obtained after coordinate rotation (from (a) to (b)) and nonlinear coordinate transformation (from (b) to (c)). Red closed curves are mutation contours defined by Eq. (14b). Green curves indicate constraint curves formed under *σ*_*y*_ = 0. The thick orange line is an evolutionary branching line detected by the branching line conditions.

## 3 Evolutionary branching in an arbitrarily distorted trait space

The above analysis in the simply distorted trait space showed that distortion of the trait space controlled by *ρ* does not affect the branching point conditions but does affect the branching line conditions. Analogous results are obtained for an arbitrarily distorted trait space of an arbitrarily higher dimension, as shown in Appendix D. In this section, for simplicity, we explain the obtained results mainly in an arbitrarily distorted two-dimensional trait space, denoted by **s** = (*x*, *y*)^T^.

### 3.1 Assumption for mutation

We generalize the assumption for the simply distorted trait space (Section 2.2) as follows (illustrated in Fig. 4a and 4b).

**Figure 4.**
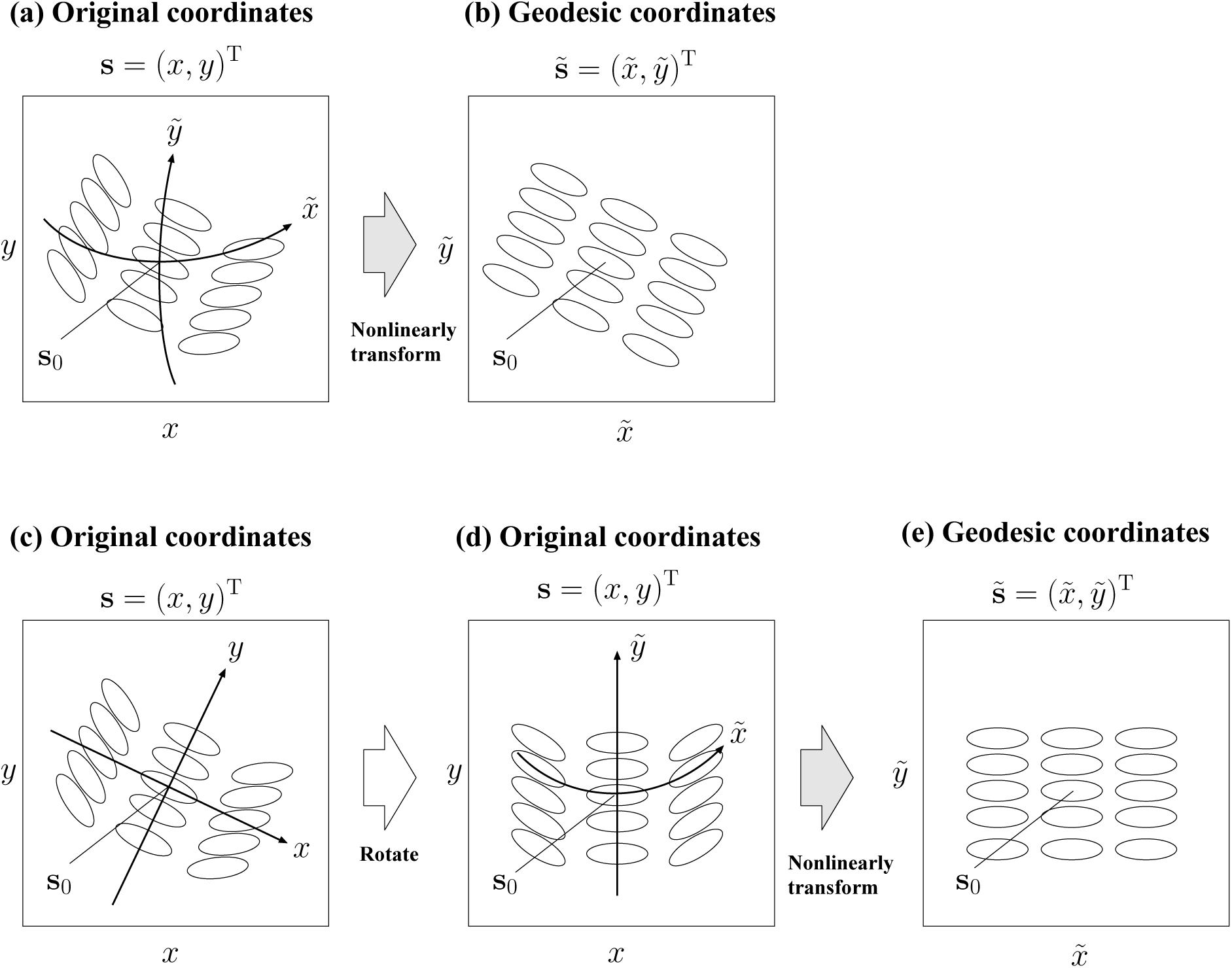
Local coordinate normalization for arbitrarily distorted trait spaces. Black ellipses are mutation ellipses, defined by Eq. (14a). A mutation ellipse indicates the mutational standard deviation in each direction from a parental phenotype located at its center.

**Geodesic-constant-mutation assumption:**

For an arbitrary point **s**_0_ = (*x*_0_, *y*_0_)^T^ in an arbitrarily distorted trait space **s** = (*x*, *y*)^T^, there exist the geodesic coordinates 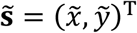 defined by

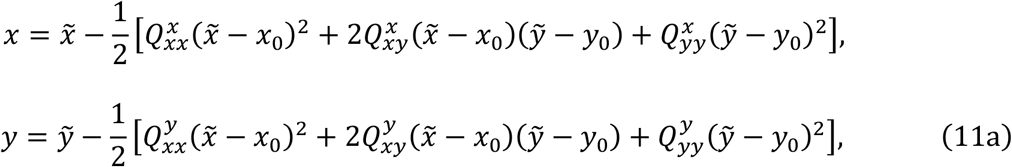

with an appropriately chosen *Q*s, such that the mutation distribution in the geodesic coordinates 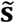 can be approximated with a constant bivariate Gaussian distribution,

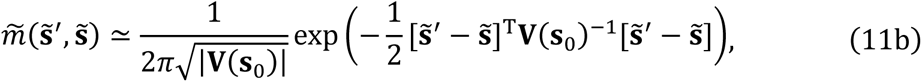

for a resident 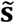 in the neighborhood of **s**_0_, satisfying

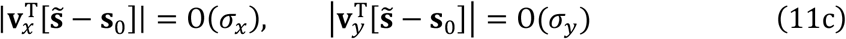

with a sufficiently small *σ*_*x*_ and *σ*_*y*_, where 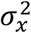 and 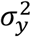 are the two eigenvalues of **V**(**s**_0_) with corresponding eigenvectors **v**_*x*_ and **v**_*y*_, respectively, and *σ*_*x*_ ≥ *σ*_*y*_ ≥ 0 is assumed without loss of generality.

The matrix **V**(**s**) in Eq. (11b) is symmetric and positive definite, referred to as a “mutational covariance matrix” or “mutational covariance”,

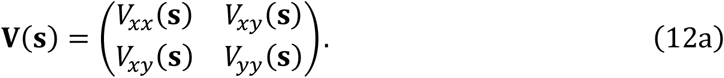

Each of the six *Q*s in Eqs. (11a) correspond to each mode of local distortion for a trait space (Fig. 5). For a given **V**(**s**), we choose 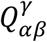 for *α*, *β*, *γ* ∈ {*x*, *y*} as

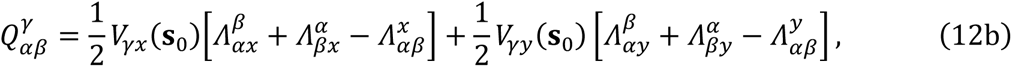

with

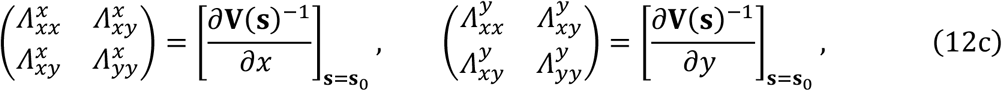

so that **V**(**s**)^−1^ has no linear dependency on 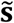 at the focal point **s**_0_ (in order to satisfy Eq. (11b)). In differential geometry, 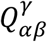 are called the Christoffel symbols of the second kind at **s**_0_ in the original coordinates **s** with respect to the metric **V**(**s**)^−1^ (see Section 3 in Hobson et al. (2006) for introduction to Christoffel symbols and geodesic coordinates). For example, in the simply distorted trait space in Section 2 (Eqs. (2)), the focal point **s**_0_ has 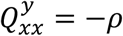 and 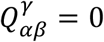 for the all other *α*, *β*, *γ* ∈ {*x*, *y*}. We refer to the inverse of the mutational covariance, **V**(**s**)^−1^, as the “mutational metric”, with which we can describe the mutational square distance from **s** to **s** + **ds** with infinitesimal **ds** = (d*x*, d*y*)^T^ as

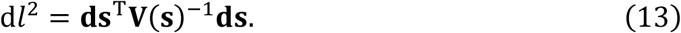

**Figure 5.**
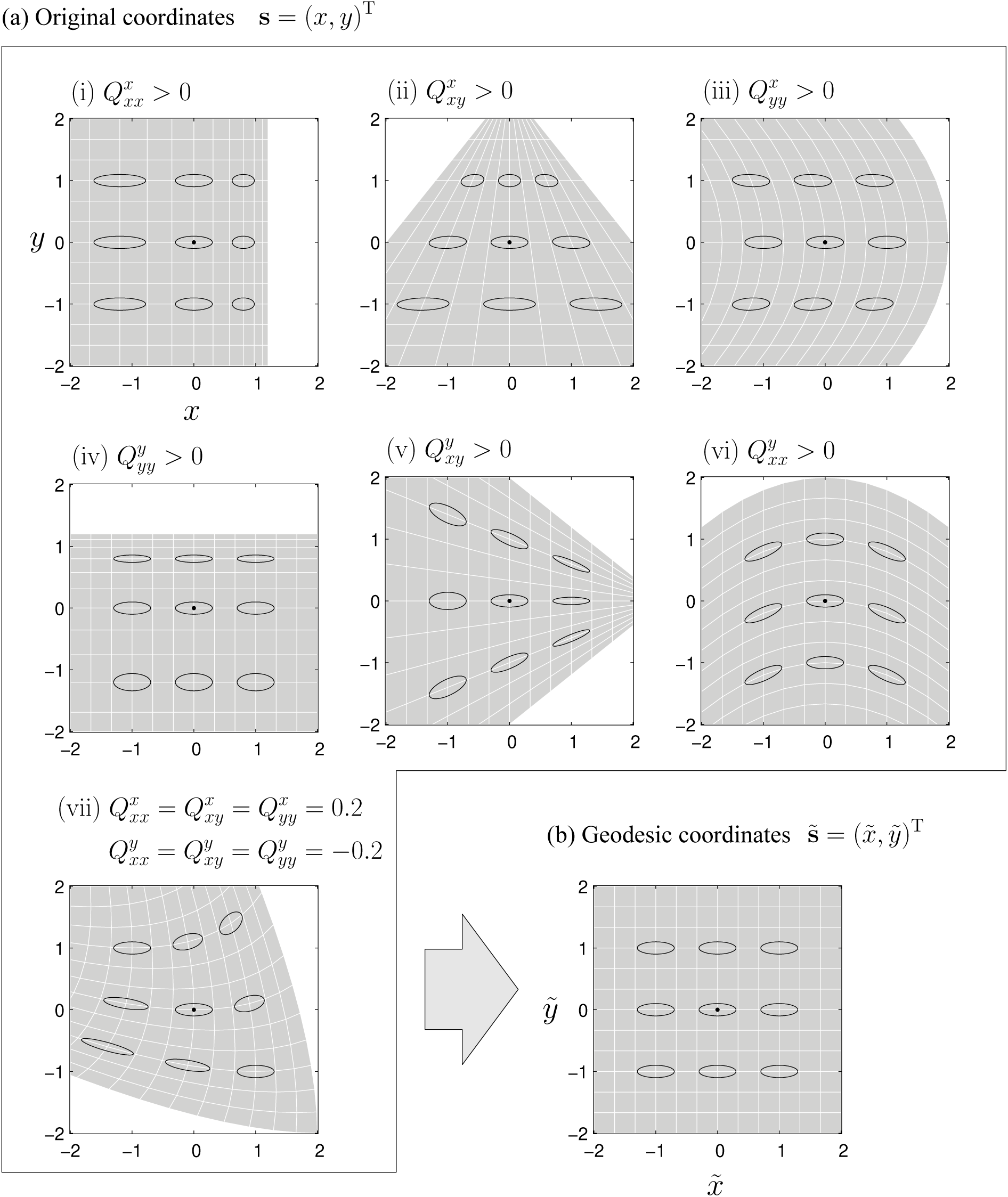
Modes of local distortion. Each of (i-vi) in panel (a) shows how each 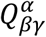 for *α*, *β*, *γ* = *x*, *y* contributes to local distortion of a trait space (only the focal 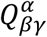 is set at 0.4, while the others are all zero). An example for all 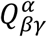s being nonzero is shown in (vii) in panel (a). All of the local distortions (i-vii) are canceled by coordinate transformation into the geodesic coordinates (Eqs. (11a)) as shown in panel (b).

Based on the mutational covariance **V**(**s**), we formally define “distorted trait spaces” as trait spaces with non-constant **V**(**s**). (This “distortion” corresponding to the first derivatives of metrics is different from the “distortion” in differential geometry defined by the second derivatives of metrics (Hobson et al., 2006).) Although the plausibility of the geodesic-constant-mutation assumption above must be examined by empirical data, this assumption provides one of the simplest frameworks that allow analytical treatment of evolutionary branching in distorted trait spaces.

In Figs. 4 and 5, the mutational covariance at each point **s**_0_ is expressed as an ellipse,

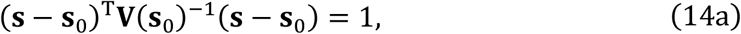

referred to as a “mutation ellipse”, which indicates the standard deviation of 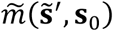 (the mutation distribution described in the geodesic coordinates) along each direction in the geodesic coordinates 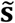 (overlaid on coordinates **s**), with its maximum and minimum given by *σ*_*x*_ and *σ*_*y*_, respectively. Expressing 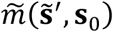 in the original coordinates **s** gives the mutation distribution *m*(**s**′, **s**_0_) in the original coordinates (see Appendix A.2 for the derivation). For *σ*_*x*_ and *σ*_*y*_ having comparable magnitudes, 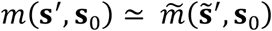 holds for sufficiently small *σ*_*x*_. In this case, the covariance matrix V_*m*_(**s**_0_) of *m*(**s**′, **s**_0_) is approximately given by the mutational covariance **V**(**s**_0_), and the mutation ellipse is approximately the same with the contour for *m*(**s**′, **s**_0_) at density level

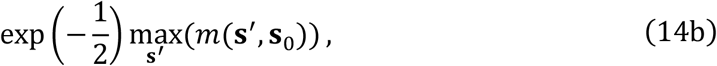

referred to as the “mutation contour” (red closed-curves in Figs. 2 and 3). On the other hand, when *σ*_*y*_ is much smaller than *σ*_*x*_, the mutation contour non-negligibly deviates from the mutation ellipse (Fig. 6). In this case, using **V**(**s**) in stead of **V**_*m*_(**s**) in Eq. (1a) may give a better description for directional evolution (see Appendix E for the details).

**Figure 6.**
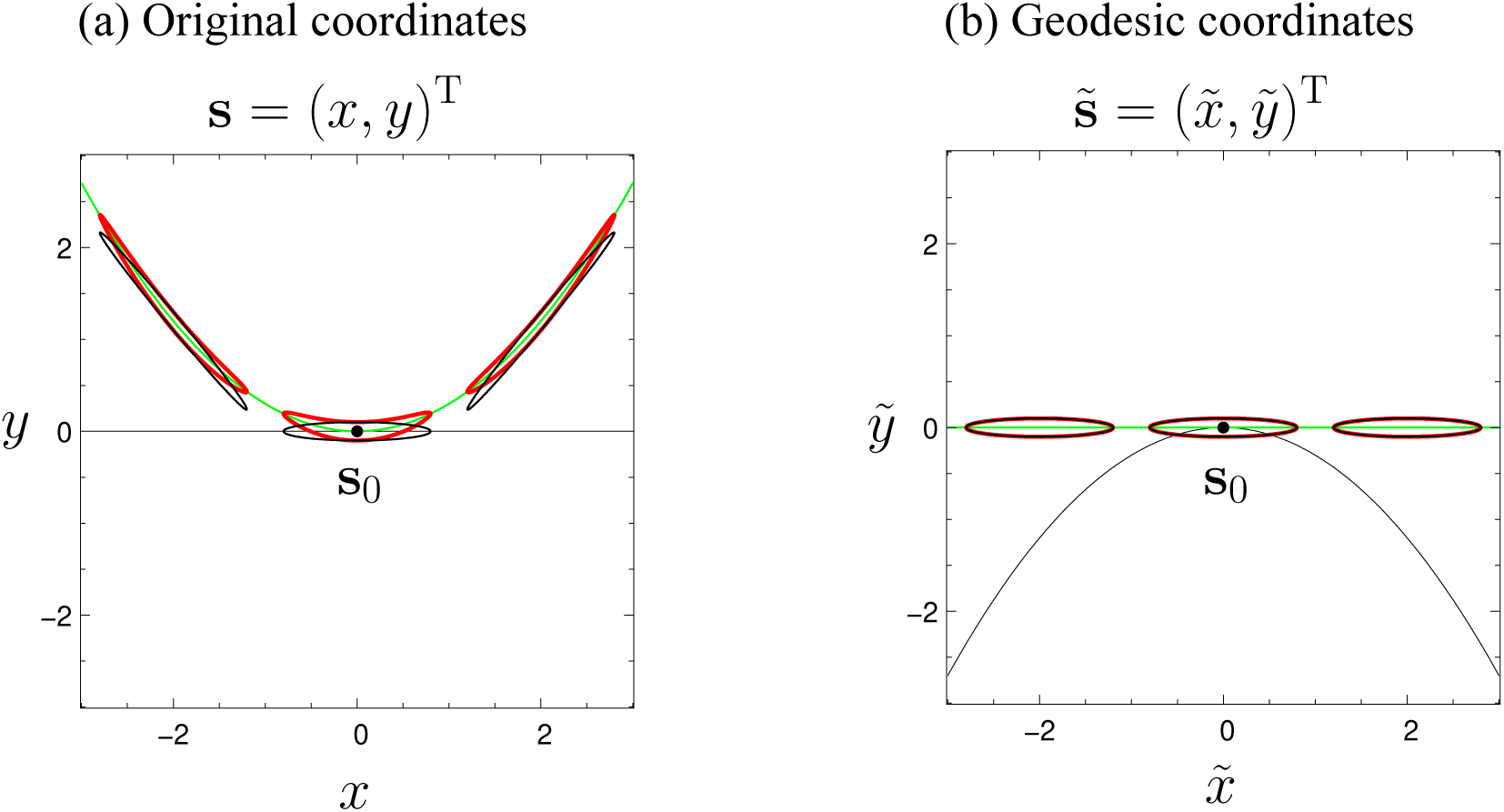
Deviation of mutation contour from mutation ellipse caused by nonlinear coordinate transformation. Black ellipses and red closed-curves are respectively mutation ellipses (Eq. (14a)) and mutation contours (Eq. (14b)). The coordinate transformation is defined by Eqs. (11a). Parameters: *σ*_*x*_ = 0.8, *σ*_*y*_ = 0.1, and 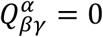 for all *α*, *β*, *γ* = *x*, *y* except 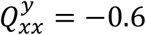.

### 3.2 Quadratic approximation of invasion fitness functions

To reduce complexity of the expressions in the subsequent analysis, without loss of generality we assume that coordinates **s** = (*x*, *y*)^T^ are first rotated so that **V**(**s**_0_) becomes a diagonal matrix expressed as

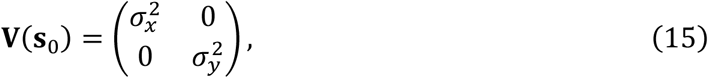

and then the geodesic coordinates 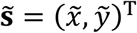 are obtained (Fig. 4c-e). In this case, Eqs. (11c) become 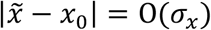 and 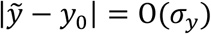. For convenience, we express Eqs. (11a) in a vector-matrix form, as

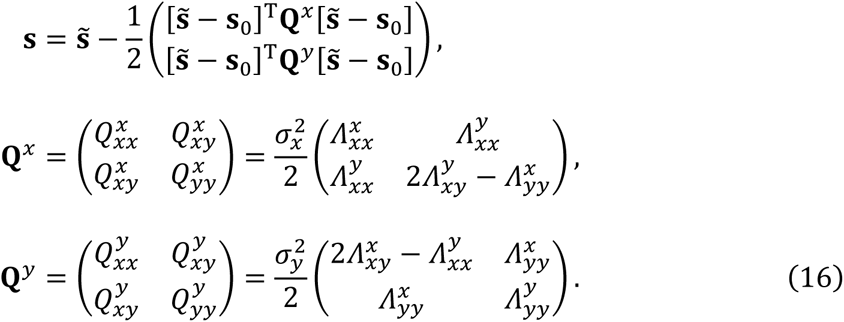

Note that **Q**^*x*^ and **Q**^*y*^ are both symmetric. We refer to **Q**^*x*^ and **Q**^*y*^ as “distortion matrices.” By substituting Eqs. (16) into the original invasion fitness function, *f*(**s**′, **s**), we derive the invasion fitness function in the geodesic coordinates 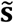, i.e., the geodesic invasion fitness,

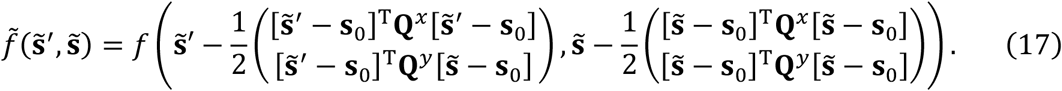

Then, we expand *f*(**s**′, **s**) in the same form with Eqs. (5), and expand 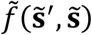 in a form similar to Eqs. (6), as

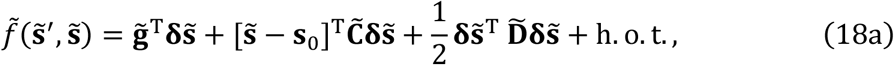

with

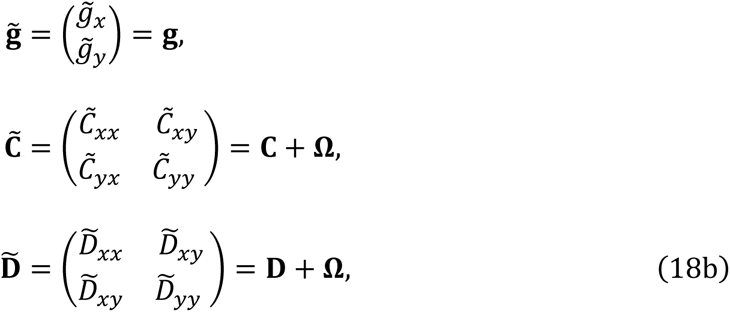

and

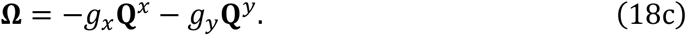

Note that Eqs. (18) are identical to Eqs. (6), except that Eq. (18c) is different from Eq. (6c).

### 3.3 Conditions for evolutionary branching points

Analogously to the branching point conditions in the simply distorted trait space (Section 2.4), we can describe conditions for a point **s**_0_ being an evolutionary branching point, as follows.

**Branching point conditions in arbitrarily distorted two-dimensional trait spaces:**

In an arbitrarily distorted trait space **s** = (*x*, *y*)^T^, a point **s**_0_ = (*x*_0_, *y*_0_)^T^ is an evolutionary branching point, if **s**_0_ satisfies the following three conditions in the corresponding geodesic coordinates 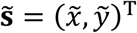 given by Eqs. (16) with Eqs. (12c) (after rotation of coordinates **s** so that Eq. (15) holds).

i. **s**_0_ is evolutionarily singular, satisfying

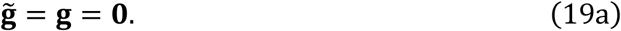
ii. **s**_0_ is strongly convergence stable, i.e., the symmetric part of

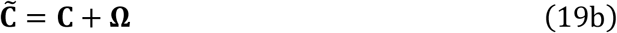

is negative definite.
iii. **s**_0_ is evolutionarily unstable, i.e., a symmetric matrix

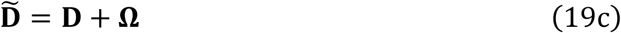

has at least one positive eigenvalue.

Here **Ω** = −*g*_*x*_**Q**^*x*^ − *g*_*y*_**Q**^*y*^, while **g**, **C**, and **D** are calculated from Eqs. (5).

Since Eq. (19a) gives **Ω** = −*g*_*x*_**Q**^*x*^ − *g*_*y*_**Q**^*y*^ = 0, we see 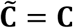 and 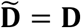. This means that the branching point conditions in the geodesic coordinates 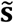 are equivalent to those in the original coordinates **s** (and in the original coordinates before the rotation). Analogous results are obtained in distorted trait spaces of arbitrary higher dimensions (Appendix D.3). Therefore, distortion of a trait space of an arbitrary dimension does not affect the branching point conditions, as long as mutation is possible in all directions.

### 3.4 Conditions for evolutionary branching lines

Analogously to the case of the simply distorted trait space in Section 2.5, when the sensitivity of the geodesic invasion fitness, 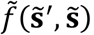, to single mutational changes of 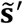 and 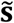 is significantly lower in 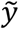 than in 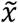, so that Eq. (9a) holds, we can simplify Eqs. (18) into

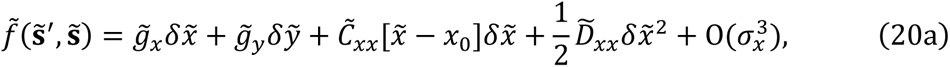

with

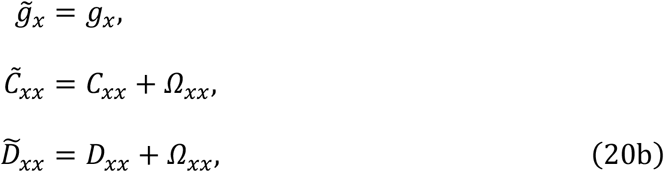

and

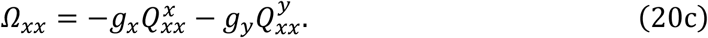

Note that Eqs. (20) are identical to Eqs. (8) except that Eq. (20c) is different from Eq. (8c). On this basis, the simplified branching line conditions for arbitrarily distorted two-dimensional trait spaces are described as follows (Appendix C.1-3).

**Branching line conditions in arbitrarily distorted two-dimensional trait spaces (simplified):**

In an arbitrarily distorted two-dimensional trait space **s** = (*x*, *y*)^T^, there exists an evolutionary branching line containing a point **s**_0_ = (*x*_0_, *y*_0_)^T^, if **s**_0_ satisfies the following four conditions in the corresponding geodesic coordinates 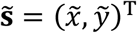 given by Eqs. (16) with Eqs. (12c) (after rotation of coordinates **s** so that Eq. (15) holds).

i. At **s**_0_ the sensitivity of the geodesic invasion fitness, 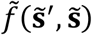, to single mutational changes of 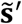 and 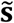 is significantly lower in 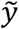 than in 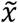, satisfying

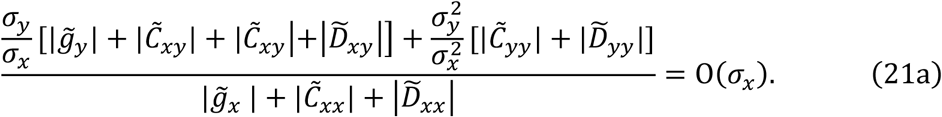
ii. **s**_0_ is evolutionarily singular along 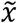, satisfying

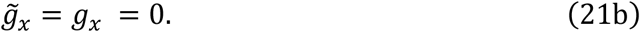
iii. **s**_0_ is convergence stable along 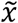, satisfying

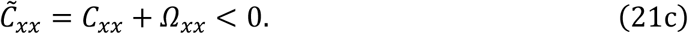
iv. **s**_0_ is sufficiently evolutionarily unstable (i.e., subject to sufficiently strong disruptive selection) along 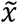, satisfying

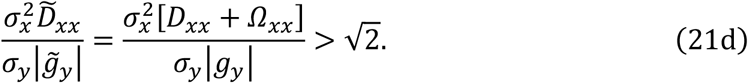

Here 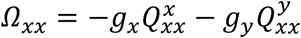, while *g*_*x*_, *g*_*y*_, *C*_*xx*_, and *D*_*xx*_ are calculated from Eqs. (5).

Note that condition (ii) *g*_*x*_ = 0 gives 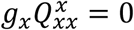, while 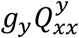 can remain nonzero in Eqs. (21c) and (21d). Thus, the distortion affects the branching line conditions through 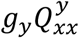, as long as the fitness gradient along the *y*-axis, *g*_*y*_, exists. Interestingly, 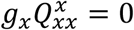 makes the above branching line conditions equivalent to the branching line conditions for the simply distorted trait space (Section 2.5), where 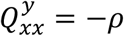. Among the six *Q*s for describing local distortion, only 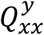 has effect on the branching line conditions, even in this general case.

When *σ*_*y*_ = 0, the evolutionary trajectory starting from **s**_0_ = (*x*_0_, *y*_0_)^T^ in coordinates 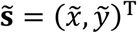 is completely restricted to the line 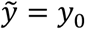, which forms a parabolic curve in the coordinates **s** = (*x*, *y*)^T^ in the neighborhood of **s**_0_,

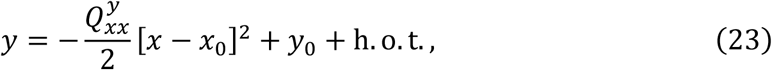

analogously to Eq. (10) in Section 2.5. In this case, condition (i) always holds, and conditions (ii-iv) become identical to the three conditions for evolutionary branching point along a constraint curve that is locally approximated in the form of Eq. (23), derived by Ito and Sasaki (2016) with an extended Lagrange multiplier method.

The branching line conditions for distorted two-dimensional trait spaces, Eqs. (21), are extended for trait spaces of arbitrary higher dimensions (Ito and Dieckmann, 2014), referred to as “candidate-branching-surface conditions” in this paper, and which are affected by the distortion in a manner analogous to the two-dimensional case here (Appendix D.4). Those conditions extend the branching point conditions along constraint curves and surfaces of arbitrary dimensions (Ito and Sasaki, 2016), for the case allowing slight mutational deviations from those curves and surfaces.

### 3.5 Conditions for evolutionary branching areas

In numerical simulations, evolutionary branching may occur before populations have reached to evolutionary branching points or lines. Consequently, the set of points where evolutionary branchings have occurred form an area or areas. To characterize such areas, Ito and Dieckmann (2012) have heuristically extended the branching line conditions into the branching area conditions, for non-distorted trait spaces. Although the branching area conditions have not been formally proved, those conditions have a good prediction performance in numerically simulated evolutionary dynamics (Ito and Dieckmann, 2012).

In this paper, we extend the branching area conditions for distorted trait spaces of two dimensions (Appendix C.5) and of arbitrary higher dimensions (Appendix D.5), by describing those conditions (for non-distorted trait spaces) in the corresponding geodesic coordinates. Analogously to the case of branching line conditions, the distortion affects the branching area conditions in trait spaces of arbitrary dimensions.

In non-distorted trait spaces, any evolutionary branching point or line is contained in an evolutionary branching area (Ito and Dieckmann, 2012). This property is kept in distorted trait spaces (Appendix C.5 and D.5).

## 4 Example

In this example, we design the trait space **s** = (*x*, *y*)^T^ by nonlinear transformation of a coordinate system having a constant Gaussian mutation distribution. This setting shows clearly how our local coordinate normalization works.

### 4.1 Ecological interaction

In trait space **s** = (*x*, *y*)^T^, we consider the two-dimensional version of the classical MacArthur-Levins resource competition model (MacArthur and Levins, 1967; Vukics et al., 2003). The growth rate of *i*th phenotype **s**_*i*_ = (*x_i_*, *y_i_*)^T^ among coexisting phenotypes **s**_1_, ⋯, **s**_*M*_ is defined by

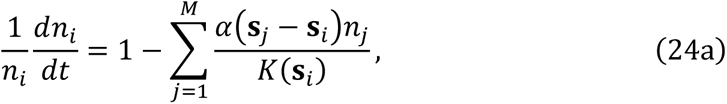

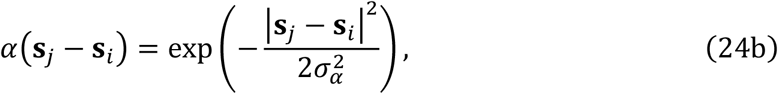

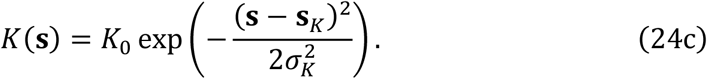

Here, *K*(**s**_i_) is the carrying capacity for phenotype **s**_*i*_, expressed with an isotropic bivariate Gaussian function with its standard deviation a_K_ and maximum *K*_0_ at **s**_*K*_ = (*x*_*K*_, *y*_*K*_)^T^. The competition kernel *α*(**s**_*j*_ − **s**_*i*_) describes the competition strength between **s**_*j*_ and **s**_*i*_, which is also an isotropic Gaussian function with its standard deviation *σ*_*α*_, i.e., the competition strength is a decreasing function about their phenotypic distance.

We assume a monomorphic population with its resident phenotype **s**, where its density n is at an equilibrium given by K(s). The invasion fitness *f*(**s**′, **s**) is defined as the per-capita growth rate of the mutant population density n′ when it is very low,

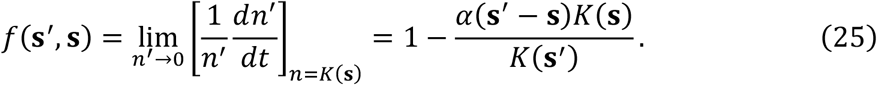

### 4.2 Mutation

To model a nontrivial but analytically tractable mutational covariance for the trait space **s** = (*x*, *y*)^T^, we assume that *x* and *y* are functions of *r* and *θ* which mutate independently, following one-dimensional Gaussian distributions with constant standard deviations *σ*_*θ*_ and *σ*_*r*_, respectively. Thus, in coordinates (*θ*, *r*)^T^, the mutation distribution is a bivariate Gaussian distribution with a constant and diagonal mutational covariance with its entries 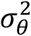 and 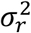 (Fig. 7b). Specifically, we define *x* and *y* as

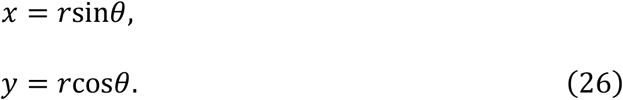

**Figure 7.**
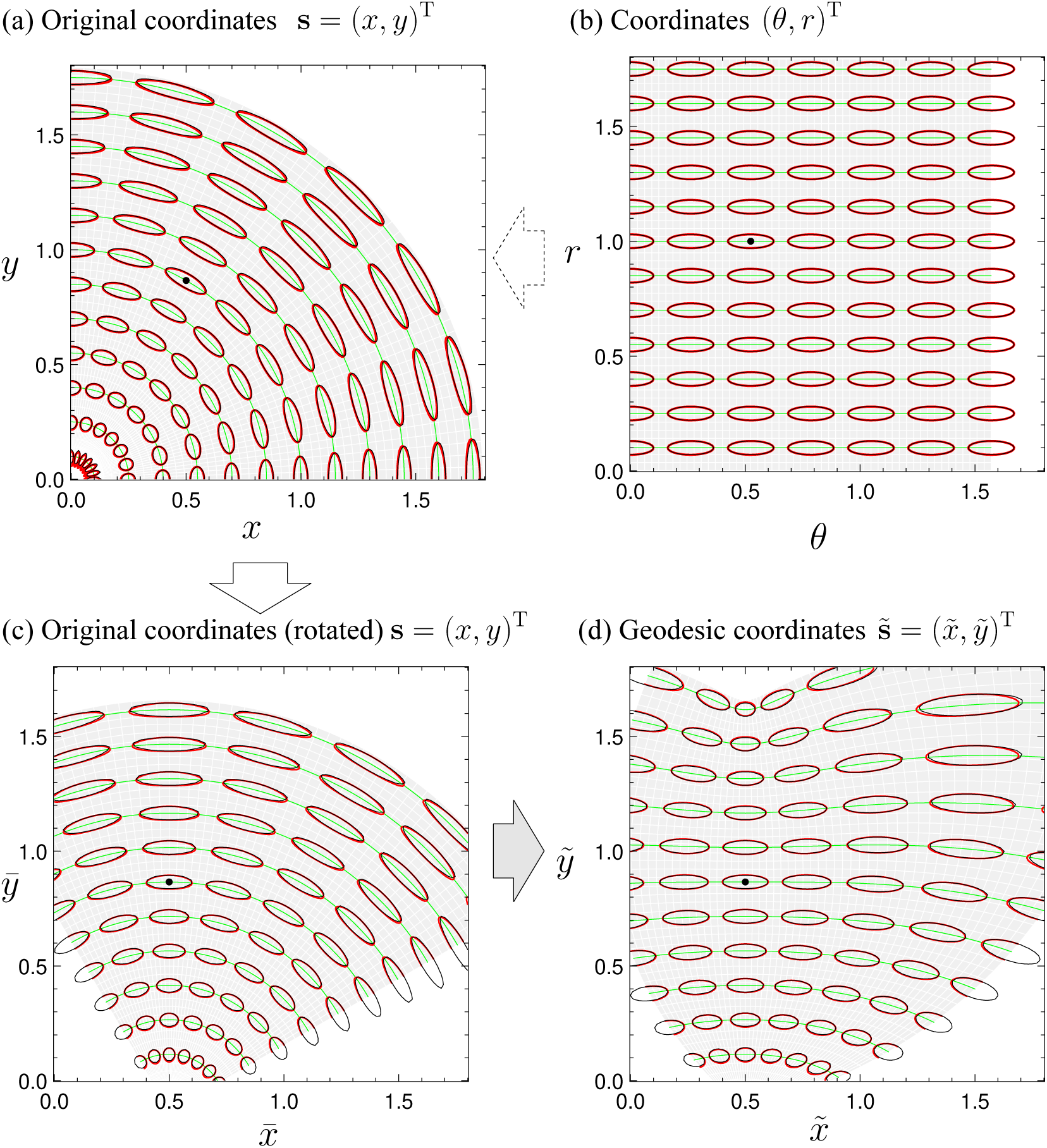
Local coordinate normalization in Example. (a) Original coordinates **s** = (*x*, *y*)^T^. (b) Non-distorted coordinates (*Θ*, *R*)^T^, nonlinear transformation of which generates the original coordinates. (c) Original coordinates after rotation, still denoted by **s** = (*x*, *y*)^T^. (d) Geodesic coordinates 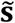. Black dots indicate a focal point **s**_0_ for examination of evolutionary branching conditions. Thick red and thin black ellipses respectively indicate mutation contours and mutation ellipses (almost identical in this figure). Green curves indicate constraint curves formed under *σ*_*r*_ = 0. Parameters: *σ*_*θ*_ = 0.2, *σ*_*r*_ = 0.03.

Eqs. (26) may be plausible when the trait space **s** = (*x*, *y*)^T^ is for predators competing for their prey animals as resources (Fig. 8), where 2*x* and *y* respectively describe the width and height of the main prey for a predator of phenotype **s** = (*x*, *y*)^T^, while *r* and *θ* respectively describe the length of predator’s jaw (or raptorial legs) and its maximum open angle. Note that both of *x* and *y* must be positive in this case.

**Figure 8.**
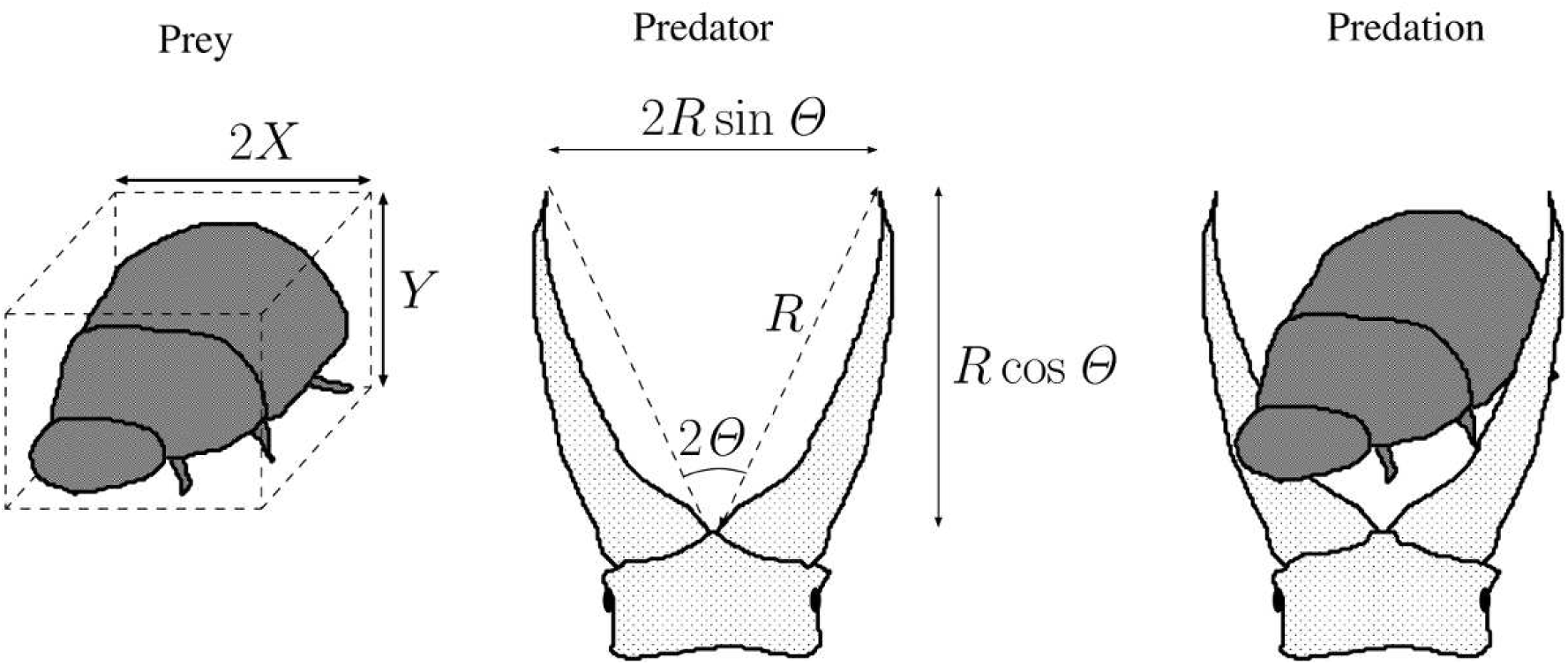
Ecological assumption for predator-prey relationship in Example.

From Eqs. (26), we can derive the mutational covariance as

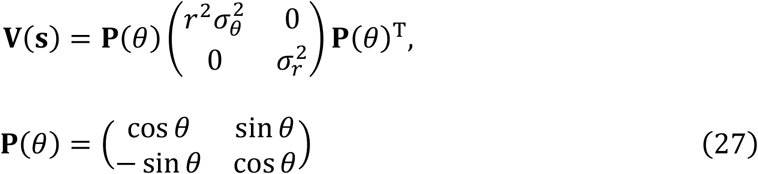

(Eq. (F.7) in Appendix F.1). After coordinate rotation about a focal point **s**_0_ = (*x*_0_, *y*_0_)^T^ = (*r*_0_ sin *θ*_0_, *r*_0_ cos *θ*_0_)^T^ so that **V**(**s**_0_) becomes diagonal, we obtain the geodesic coordinates 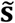 (Fig. 7d) with

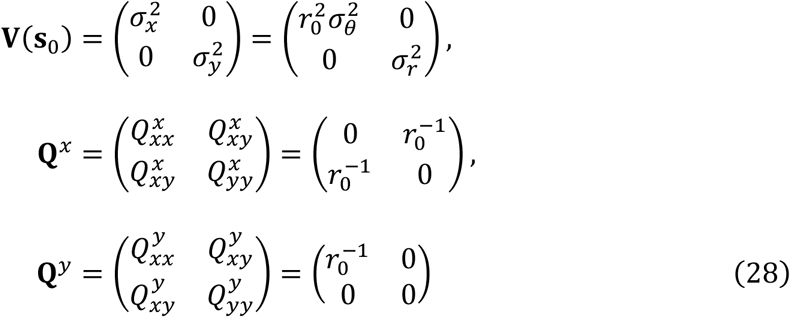

(Appendix F.2). Note that the constant Gaussian mutation distribution in coordinates (*θ*, *r*)^T^ (Fig. 7b) is locally recovered around the focal point **s**_0_ in the geodesic coordinates 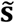 (Fig. 7d). In this special example, the non-distorted coordinates (*θ*, *r*)^T^ allows application of the evolutionary branching conditions for non-distorted trait spaces. (As shown in Appendix F. 6, the branching point conditions and branching line conditions derived in the non-distorted coordinates (*r*, θ)^T^ are identical to those in the geodesic coordinates 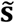.) However, obtaining such coordinates is usually impossible for a given mutational covariance **V**(**s**). On the other hand, the local coordinate normalization by obtaining the geodesic coordinates 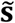 by the local coordinate normalization is possible in many cases.

### 4.3 Branching point conditions

From Eq. (25), we derive the fitness gradient, fitness-gradient variability, and fitness curvature at the focal point **s**_0_ in the original coordinates (after rotation, Fig. 7c), as

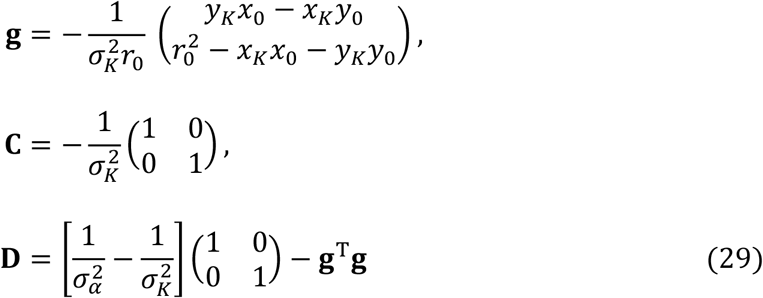

(Appendix F.3). As shown in Section 3.3, the branching point conditions (Eqs. 19) are not affected by the distortion. Thus, we can directly examine those conditions in the original coordinates **s**. Consequently, a necessary and sufficient condition for the existence of an evolutionary branching point is given by *σ*_*α*_ < *σ*_*K*_. When *σ*_*α*_ < *σ*_*K*_, an evolutionary branching point exists at the peak point of the carrying capacity, **s**_*K*_ (Appendix F.4), as already derived in Vukics et al. (2003) for non-distorted trait spaces. Conversely, when *σ*_*α*_ > *σ*_*K*_, the point **s**_*K*_ is locally evolutionarily stable as well as strongly convergence stable, in which case **s**_*K*_ is not an evolutionary branching point.

### 4.4 Branching line conditions

The branching line conditions (Eqs. (21)) are examined in this model by substituting Eqs. (28) and (29) into Eqs. (21). As shown in Appendix F.5, for a *σ*_*r*_ sufficiently smaller than *σ*_*θ*_, there exists an evolutionary branching line along the line passing through the origin and the peak point **s**_*K*_ = (*x*_*K*_, *y*_*K*_)^T^ of the carrying capacity, expressed in the original coordinates before the rotation, as

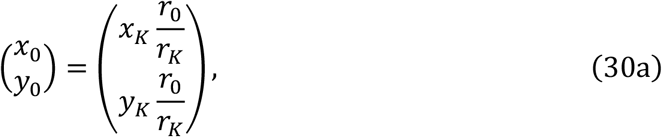

with 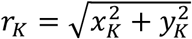 and a positive parameter *r*_0_, where the range of *r*_0_ is given by

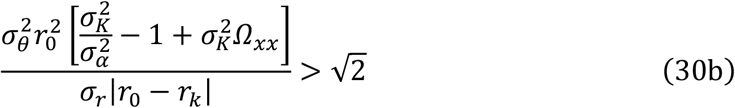

with

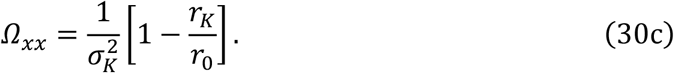

Note that this branching line exists even under *σ*_*α*_ > *σ*_*K*_, in which case there exists no branching point. Moreover, the distortion effect *Ω*_*xx*_ enables the existence of this branching line, because Eq. (30b) is never satisfied for *Ω*_*xx*_ = 0 under *σ*_*α*_ > *σ*_*K*_.

### 4.5 Numerical analysis

Figure 9 shows evolutionary dynamics simulated numerically as trait substitution sequences (Ito and Dieckmann, 2014) starting from various initial phenotypes, under *σ*_*r*_ = *σ*_*θ*_ (see Appendix G for the simulation algorithm). This simulation assumes *σ*_*α*_ > *σ*_*K*_, i.e., the unique evolutionary singular point **s**_*K*_ is convergence stable but not an evolutionary branching point. As predicted, all evolutionary trajectories converge to **s**_*K*_, but evolutionary branching does not occur. Even in this case, a branching line can exist when *σ*_*r*_ is much smaller than *σ*_*θ*_ (Fig. 10a), inducing evolutionary branching (Fig. 10c-e). The area of occurrence of evolutionary branchings is well characterized by the branching area (Fig. 10b).

**Figure 9.**
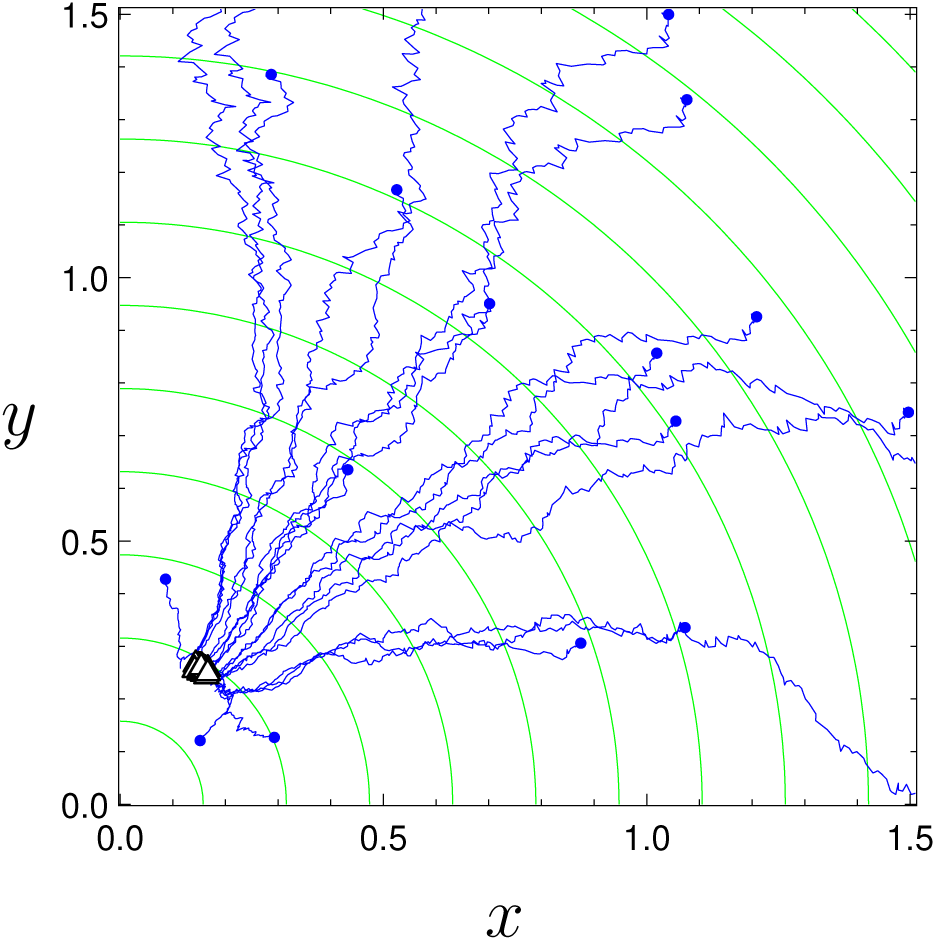
Numerically calculated evolutionary trajectories in Example, without significant anisotropy in mutation. From each of randomly chosen 25 initial phenotypes (small blue dots) within 0 ≤ *θ* ≤ *π*/2 and 0.1 ≤ *r* ≤ 1.5, evolutionary trajectories was calculated for 10^9^ generations, as a trait substitution sequence assuming asexual reproduction (blue curves) (see Appendix G for the simulation algorithm). White triangles bordered with black indicate the final resident phenotypes that have not brought about evolutionary branching. Neither evolutionary branching line (Section 3.4) nor area (Section 3.5) was found (condition (i) in the branching line condition is examined by replacing “= O(*σ*_*x*_)” with 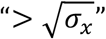 in the right hand side of Eq. (21a)). Parameters: *x*_*K*_ = 0.15, 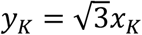, K_0_ = 1 × 10^6^, *σ*_*K*_ = 0.7, *σ*_*α*_ = 0.75, *μ* = 1 × 10^−5^ (mutation rate per birth event), and *σ*_*θ*_ = *σ*_*r*_ = 5 × 10^−3^.

**Figure 10.**
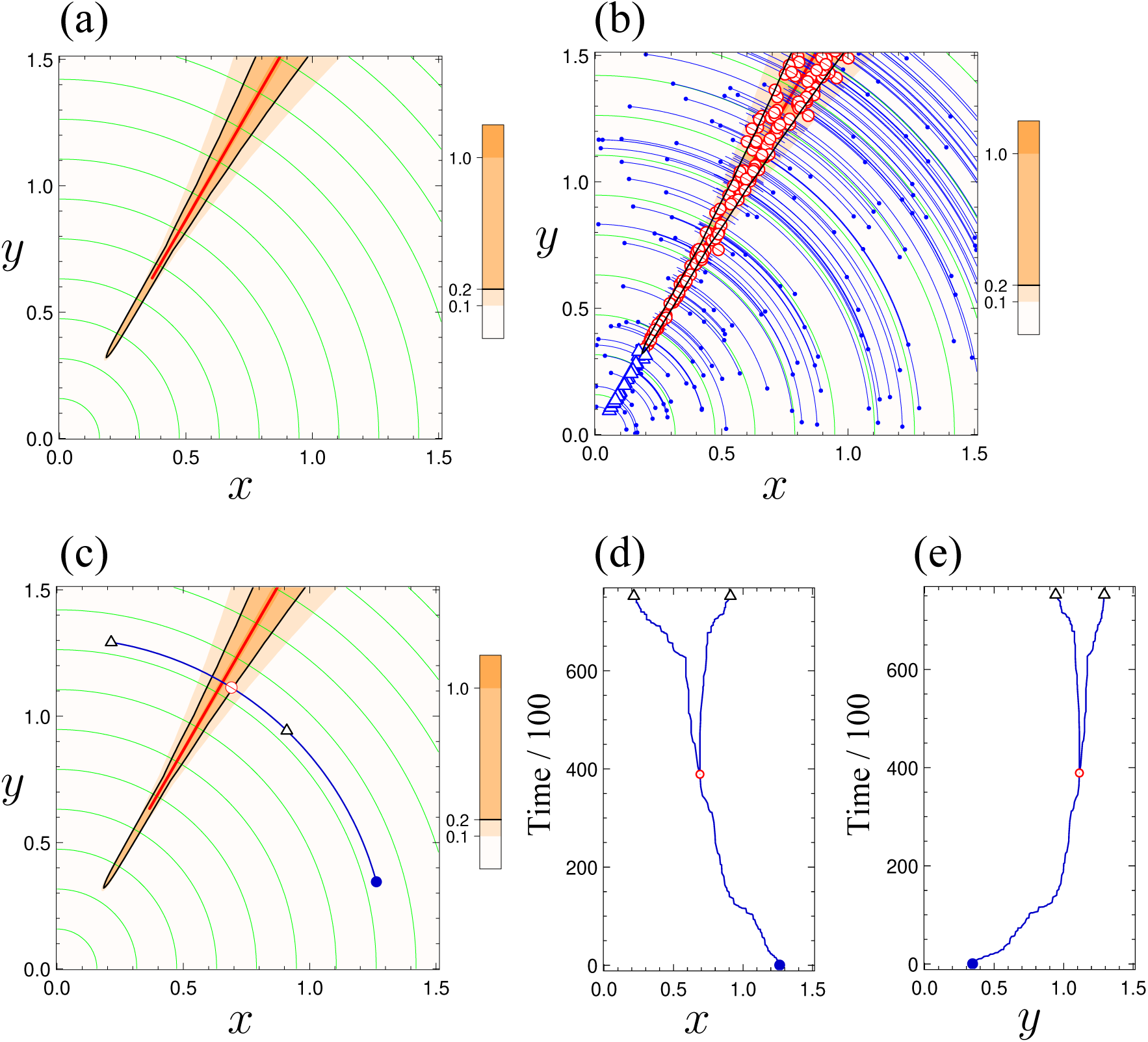
Comparison of evolutionary branching lines and areas with numerically calculated evolutionary trajectories in Example, with significant anisotropy in mutation. In panel (a), an evolutionary branching line (Section 3.4) is indicated with a red line. An evolutionary branching area (Section 3.5) is indicated with an orange area bordered by black curve (values of the color bar indicate the values for *β* in Eqs. (C.7) in Appendix C.5). The green curves indicate constraint curves formed under *σ*_*r*_ = 0. Panel (b) shows 50 evolutionary trajectories numerically calculated as trait substitution sequences (blue curves) for 10^9^ generations (Appendix G), with initial phenotypes (small blue dots) randomly chosen within 0 ≤ *θ* ≤ *π*/2 and 0.1 ≤ *r* ≤ 1.5. White circles bordered with red indicate occurrence of evolutionary branching there, while white triangles bordered with blue indicate the final resident phenotypes that have not brought about evolutionary branching. Panels (c-e) show an example evolutionary trajectory (blue curves). The initial state, first branching, and the state at the end of simulation are indicated with the blue filled circle, white circle bordered with red, white triangle bordered with black, respectively. The time unit in panels (d-e) is generation. Parameters: *σ*_*θ*_ = 5 × 10^−3^, *σ*_*r*_ = 1 × 10^−5^, and other parameters are the same as in Fig.9.

Therefore, both of analytical and numerical results in this example accord with the general result derived in Section 3 that distortion of a trait space affects evolutionary branching when mutatability have significant magnitude differences among directions, through the branching line conditions and branching area conditions.

## 5 Discussion

### 5.1 General discussion

Biological communities are thought to have been evolving in trait spaces that are not only multi-dimensional but also distorted, in a sense that mutational covariance matrices depend on the parental phenotypes of mutants. For efficient analysis of adaptive evolutionary diversification in distorted trait spaces, we made an assumption for mutation such that an appropriate local nonlinear coordinate transformation allows approximation of the mutation distribution with a locally constant Gaussian distribution, and then we applied conventional conditions for evolutionary branching points (Metz et al., 1996; Geritz et al., 1997), lines (Ito and Dieckmann, 2014) and areas (Ito and Dieckmann, 2012). Consequently, we have shown that the distortion does not affect branching point conditions but do affect branching line conditions and area conditions, in two-dimensional trait spaces. Analogous results have been obtained in trait spaces of arbitrary higher dimensions (Appendix D). Our method provides an extension tool of adaptive dynamics theory for distorted trait spaces. Our assumption for mutation and coordinate normalization described in Subsection 3.1 may be useful in other theories for evolution as well, including quantitative genetics.

### 5.2 Assumption for mutation and evolutionary constraints

Although the plausibility of our assumption for the geodesic-constant-mutation in Section 3.1 must be examined by empirical data, our assumption provides one of the simplest frameworks that allow analytical treatment of evolutionary branching in distorted trait spaces. The biological plausibility of our assumption may be examined by using it as a statistical fitting function for empirical data about mutation distributions, and checking its fitting performance. At least, our assumption would be closer to the reality than assuming a constant mutation distribution over a trait space.

An advantage of our assumption is that evolutionary dynamics along constraint curves or surfaces (of arbitrary dimensions) can be described by setting zeros for some eigenvalues of the mutational covariance matrix. The obtained evolutionary branching conditions are identical to those derived by Ito and Sasaki (2016) with an extended Lagrange multiplier method. (The obtained conditions are also mathematically equivalent to deMazancourt and Dieckmann (2004) when constraints are one-dimensional curves in two-dimensional trait spaces, and to Kisdi (2015) when constraints are one-dimensional curves in trait spaces of arbitrary dimensions.) Note that in reality all constraint curves are incomplete in a sense that mutational deviations from the curves must not be impossible, although their magnitudes may be very small and/or their likelihoods may be very low. Therefore, constraint curves themselves can change evolutionarily. Under our assumption for mutation in this paper, such a situation is easily expressed by assuming very small but nonzero values for some eigenvalues of the mutational covariance matrix, which gives evolutionary branching conditions along incomplete constraints, in the form of the branching line conditions for two-dimensional trait spaces (Sections 3.4) and the candidate-branching-surface conditions for arbitrary higher-dimensional trait spaces (Appendix D.4).

### 5.3 Relationship between evolutionary branching points and lines

If a focal point **s**_0_ is an evolutionary branching point with positive 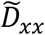 (Section 3.3), the point also satisfies the branching line conditions for sufficiently small *σ*_*y*_ (Section 3.4), which allows the coexistence of the branching point and a branching line containing the point, like as Fig.2 in Ito and Dieckmann (2012) for a non-distorted two-dimensional trait space. On the other hand, if the focal point is an evolutionary branching point with negative 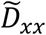, the branching line conditions are not satisfied by any small *σ*_*y*_. In this case, *σ*_*y*_ → 0 makes the branching point just vanish. On the other hand, depending on the invasion fitness function, a sufficiently small *σ*_*y*_ allows existence of a branching line containing no branching point, as shown in Fig. 10, like as Figs. 4 in Ito and Dieckmann (2012) for non-distorted two-dimensional trait spaces. Thus, the relationship between branching points and branching lines is complex, requiring further analyses.

### 5.4 Comparison with population genetic theory for distorted trait spaces

Rice (2002) developed a general population genetic theory for the evolution of developmental interactions, in the framework of quantitative genetics. This theory can analyze evolutionary dynamics in distorted trait spaces from the perspective of developmental interactions, while its focal time span is different from our method. The theory by Rice (2002) seems good for analyzing short-term evolution with explicit description of the dynamics of standing genetic variations, while our method is good for analyzing long-term directional evolution and evolutionary diversification with simplification of the genetic structure.

## Appendix A: Mutation distributions in original coordinates

### A.1. Simply distorted trait space

By using Eqs. (3) in the main text, the mutation distribution *m*(**s**′, **s**) can be expressed as

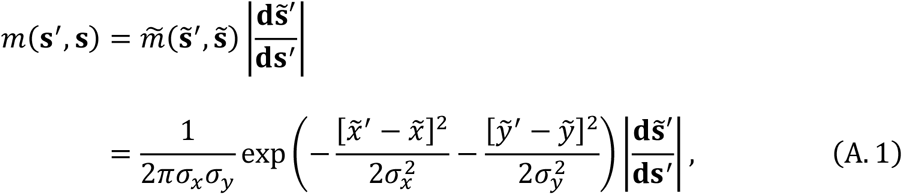

where the expansion or diminishing rate of area element due to the coordinate transformation is described by 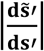, which is the determinant of a Jacobian matrix 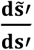. By expressing Eqs. (2) with respect to mutant **s**′,

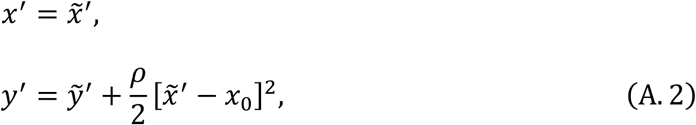

we see

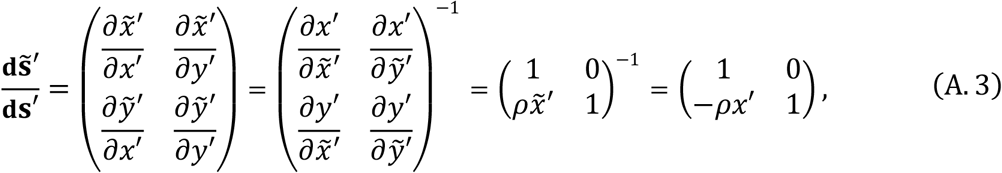

and thus

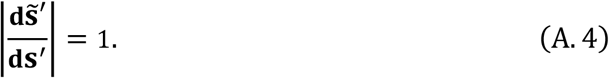

In addition, from Eqs. (2) we see

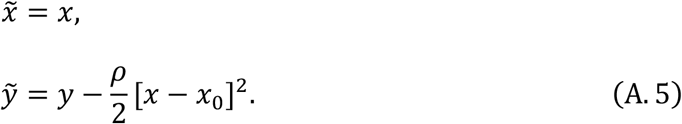

Substituting Eqs. (A.4) and (A.5) into Eq. (A.1) gives

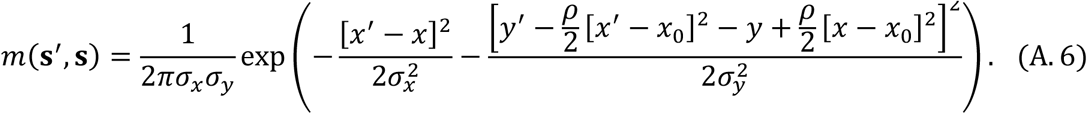

To see the deviation of *m*(**s**′, **s**) from 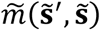, we express *δx* = *x*′ − *x*, *δy* = *y*′ − *y* as

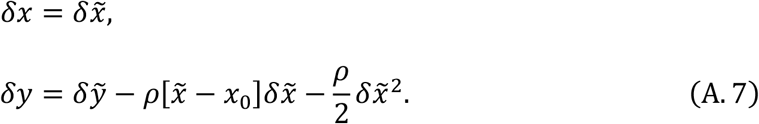

with 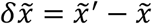, and 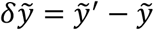. Note that 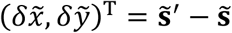 follows 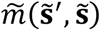, which is a constant Gaussian distribution, Eqs. (3). If *σ*_*y*_ has a similar magnitude to *σ*_*x*_, i.e., both of 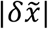 and 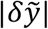 are expected to be of order 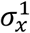, then 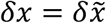 and 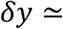 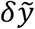 hold for a resident in the neighborhood of the focal point satisfying 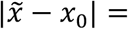 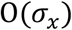. In this case, the deviation of *m*(**s**′, **s**) from 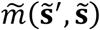 is negligible. On the other hand, if *σ*_*y*_ is much smaller than *σ*_*x*_ so that 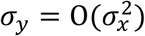 i.e., 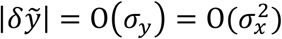 is expected, then 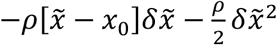 is not negligible, and thus 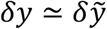 does not hold. In this case, the deviation of *m*(**s**′, **s**) from 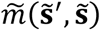 is not negligible, even under an extremely small *σ*_*x*_.

### A.2. Arbitrarily distorted trait space

For an arbitrarily distorted two-dimensional trait space, from Eqs. (11) we express the mutation distribution *m*(**s**′, **s**) as

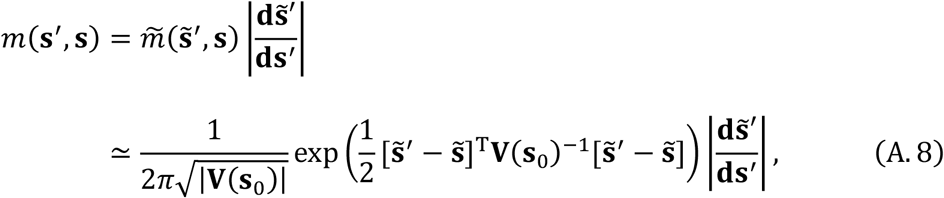

where 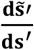 is expressed by using Eq. (16) as

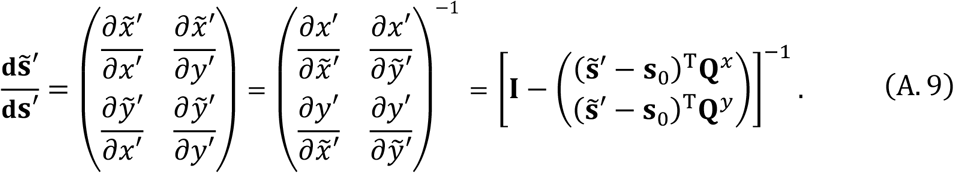

From Eqs. (16), we can express 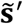 and 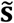 as

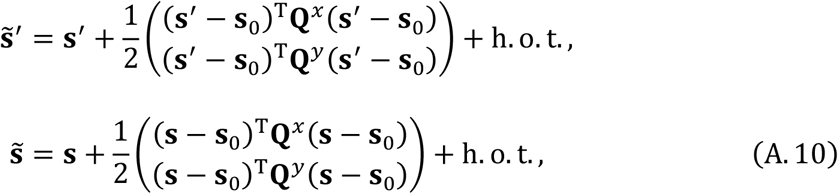

from which we express 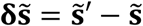 as

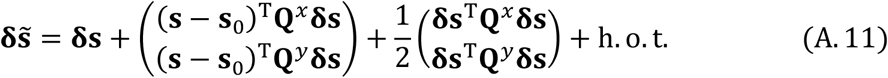

with **δs** = **s**′ − **s**. Substituting Eqs. (A.9) and (A.11) into Eq. (A.8) approximately gives an explicit form for *m*(**s**′, **s**), which is used for plotting mutation contours in Fig. 7.

To see the deviation of *m*(**s**′, **s**) from 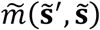, we express *δx* = *x*′ − *x*, *δy* = *y*′ − *y* as

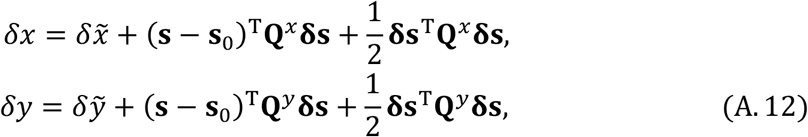

with 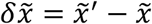, and 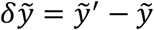. Then analogously to Section A.1 above, if *σ*_*y*_ has a similar magnitude to *σ*_*x*_, the deviation of *m*(**s**′, **s**) from 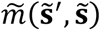 is negligible. On the other hand, if *σ*_*y*_ is much smaller than *σ*_*x*_ so that 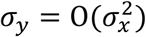, the deviation of *m*(**s**′, **s**) from 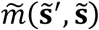 is not negligible, even under an extremely small *σ*_*x*_.

## Appendix B: Quadratic approximation of invasion fitness functions

Following Ito and Dieckmann (2014), we derive an approximate quadratic form of *f*(**s**′, **s**), as follows. We assume **s**_0_ = 0 without loss of generality. We expand *f*(**s**′, **s**) around **s**_0_ = 0 with respect to **s**′ and **s** as

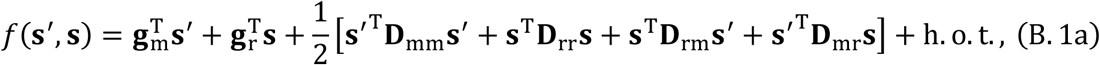

with

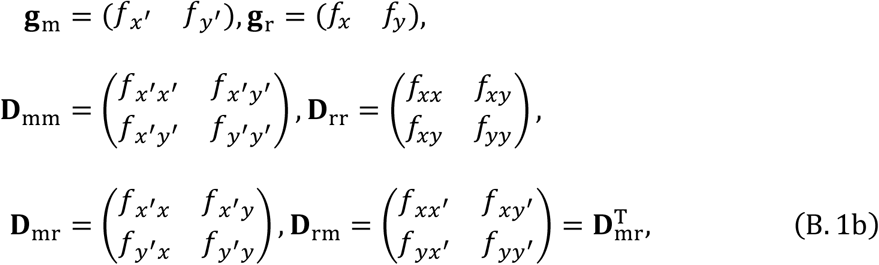

where the subscripts ‘m’ and ‘r’ refer to mutants and residents, respectively, and where *f*_*α*_ = *∂f*(**s**′, **s**)/*∂α* for *α* = *x*′, *y*′, *x*, *y* and *f*_*αβ*_ = *∂*^2^*f*(**s**′, **s**)/*∂α∂β* for *α*, *β* = *x*′, *y*′, *x*, *y* denote the first and second derivatives of *f*(**s**′, **s**), respectively, evaluated at **s**′ = **s** = **s**_0_. Since *f*(**s**, **s**) = 0 always holds for any **s** by definition, we see from Eq. (B.1a) that

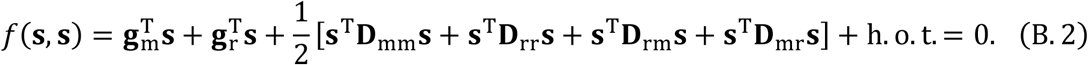

Subtracting Eq. (B.2) from Eq. (B.1a) gives

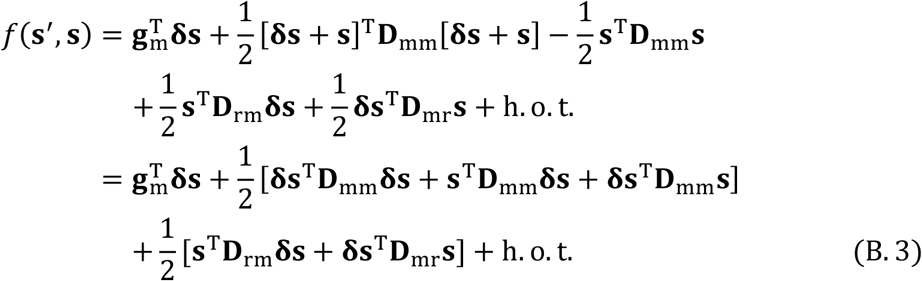

with **δs** = **s**′ − **s**. By using **δs**^T^D_mm_**s** = [**δs**^T^D_mm_**s**]^T^ = **s**^T^D_mm_**δs** = **s**^T^D_mm_**δs** and **δs**^T^D_mr_**s** = [**δs**^T^D_mr_**s**]^T^ = **s**^T^D^T^ **δs** = **s**^T^D_rm_**δs**, we further transform Eq. (B.3) into

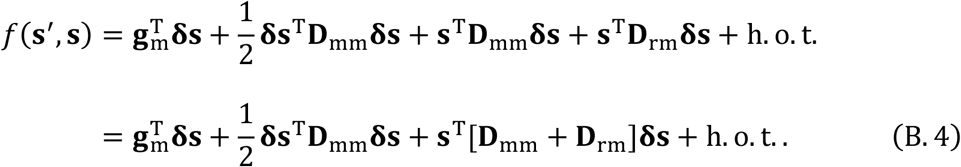

Analogously, for **s**_0_ ≠ **0**, we get

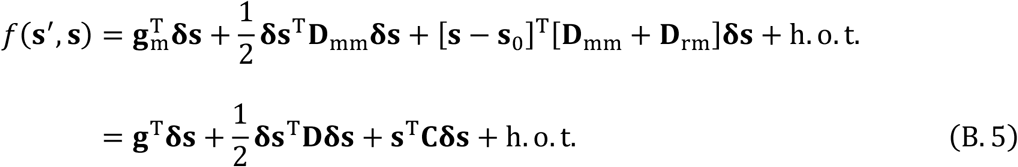

with **g** = **g**_m_, **D** = **D**_mm_, and **C** = **D**_mm_ + **D**_rm_.

## Appendix C: Branching line conditions and area conditions in arbitrarily distorted two-dimensional trait spaces

### C.1. Preparation

To apply the original branching line conditions (Ito and Dieckmann, 2014) to a distorted trait space, we transform the geodesic coordinates 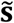 for a focal point **s**_0_, given by Eqs. (12c), (15) and (16), into new coordinates 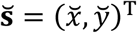, so that the mutational covariance becomes *σ*_*x*_**I**, i.e., the mutation is isotropic with its standard deviation *σ*_*x*_. Specifically, we define coordinates 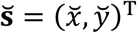 by

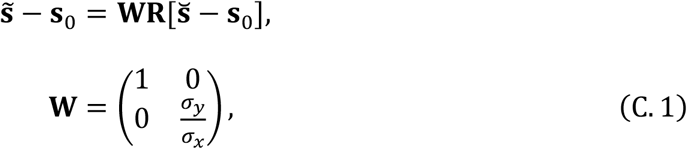

where **R** is a rotation matrix for further adjustment, which is used for description of the original branching line conditions and area conditions. Substituting Eqs. (C.1) into the geodesic invasion fitness, Eqs. (18) in the main text, gives the invasion fitness in coordinates 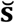,

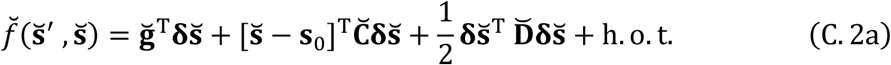

with

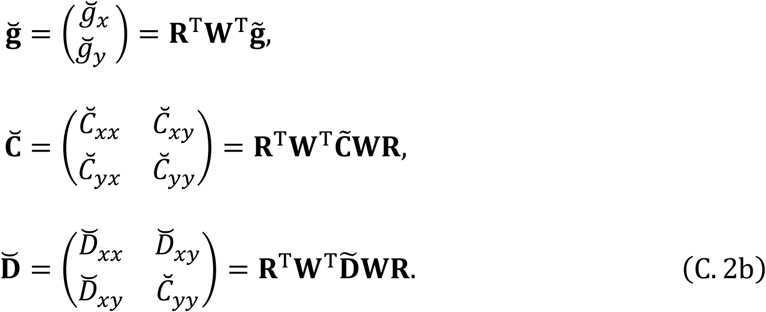

According to Eqs. (11b) and (11c) in the geodesic-constant mutation assumption, the mutation distribution 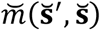 in coordinates 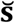 can be approximated with a constant isotropic Gaussian distribution with its standard deviation *σ*_*x*_,

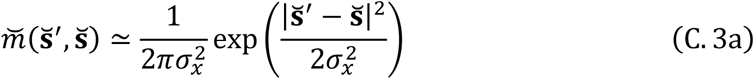

for a resident 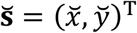 satisfying

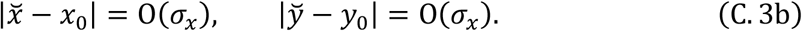

Thus, if the focal point **s**_0_ satisfies the branching line conditions below (or the branching point conditions), we expect that evolutionary branching successfully proceed (from an initial resident 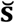 satisfying 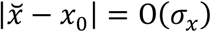 and 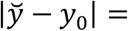 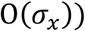, as long as distances of coexisting residents to the focal point **s**_0_ are all O(*σ*_*x*_), so that the mutation distributions for those residents still can be approximated with a constant and isotropic Gaussian distribution with its standard deviation *σ*_*x*_, and that the quadratic approximation of the invasion fitness function is valid (Ito and Dieckmann, 2014).

### C.2. Original branching line conditions

In coordinates 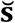 defined in Eqs. (C.1), we describe the original branching line conditions (Ito and Dieckmann, 2014), as follows.

Branching line conditions in arbitrarily distorted two-dimensional trait spaces (original)

In an arbitrarily distorted two-dimensional trait space **s** = (*x*, *y*)^T^, there exists an evolutionary branching line containing a point **s**_0_ = (*x*_0_, *y*_0_)^T^, if **s**_0_ satisfies the following four conditions in the corresponding coordinates 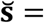 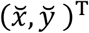 given by Eqs. (C.1), (12c), and (16) (after rotation of coordinates **s** so that Eq. (15) holds), with an appropriate choice of **R**.

i. At **s**_0_ the sensitivity of 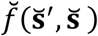 to single mutational changes of 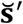 and 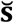 are significantly lower in 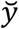 than in 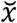, satisfying

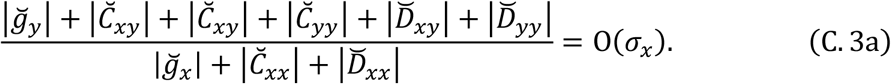
ii. **s**_0_ is evolutionarily singular along 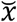, satisfying

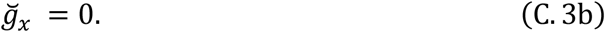
iii. **s**_0_ is convergence stable along 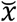, satisfying

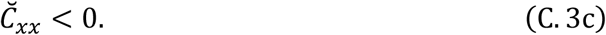
iv. **s**_0_ is sufficiently evolutionarily unstable (i.e., subject to sufficiently strong disruptive selection) along 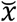, satisfying

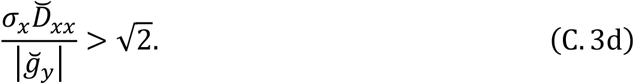

Note that condition (i) allows simplification of Eq. (C.2a) into

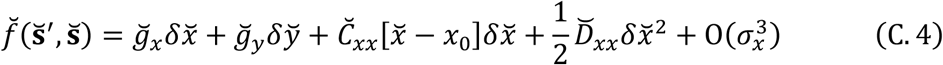

(Ito and Dieckmann, 2014).

### C.3. Simplified branching line conditions

When 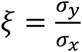 is very small so that 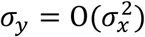, an appropriate **R** for the original branching line conditions is approximately given by **I** (Ito and Dieckmann, 2014). Assuming **R** = **I** transforms Eqs. (C.2b) into

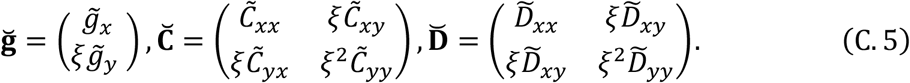

Substituting Eqs. (C.5) into Eqs. (C.3) gives the simplified branching line conditions in Section 3.4.

### C.4. Original branching area conditions

In coordinates 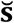 defined by Eqs. (C.1), we can apply the original branching area conditions (Ito and Dieckmann, 2012), as described below.

Branching area conditions in arbitrarily distorted two-dimensional trait spaces (original)

In an arbitrarily distorted two-dimensional trait space **s** = (*x*, *y*)^T^, there exists an evolutionary branching area containing a point **s**_0_, if **s**_0_ satisfies the following two conditions in the corresponding coordinates 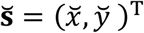 given by Eqs. (C.1), (12c), and (16) (after rotation of coordinates **s** so that Eq. (15) holds), where **R** is chosen so that 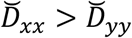 and 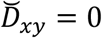 hold.

i. **s**_0_ satisfies

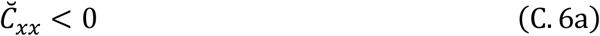

(i.e., convergence stable along 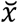 when 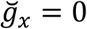).
ii. **s**_0_ satisfies

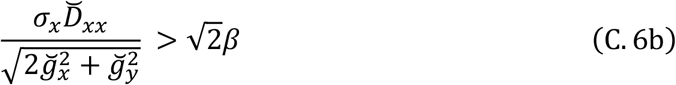

(i.e., sufficiently evolutionarily unstable along 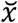 when 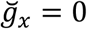).

The *β* is a positive constant to prevent condition (ii) from being too conservative. Since 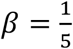 has shown a good prediction performance in Ito and Dieckmann (2012), 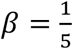 is used in this paper as well.

### C.5. Simplified branching area conditions

When *σ*_*y*_ ≪ *σ*_*x*_ holds, **R** for attaining 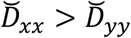 and 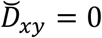 in the branching area conditions above is approximately given by **I**. Then **R** = **I** transforms Eqs. (C.2b) into Eqs. (C.5). Substituting Eqs. (C.5) into Eqs. (C.6) gives the simplified branching area conditions, as described below.

Branching area conditions in arbitrarily distorted two-dimensional trait spaces (simplified):

In an arbitrary distorted two-dimensional trait space **s** = (*x*, *y*)^T^, there exists an evolutionary branching area containing a point **s**_0_, if **s**_0_ satisfies the following two conditions in the corresponding geodesic coordinates 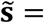 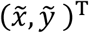 given by Eqs. (12c), and (16) (after rotation of coordinates **s** so that Eq. (15) holds), under *σ*_*y*_ ≪ *σ*_*x*_.

i. **s**_0_ satisfies

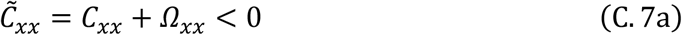

(i.e., convergence stable along 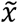 when *g*_*x*_ = 0).
ii. **s**_0_ satisfies

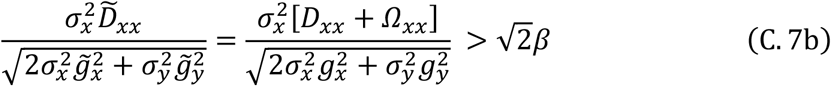

(i.e., sufficiently evolutionarily unstable along 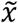 when *g*_*x*_ = 0), where 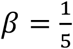.

Under *σ*_*y*_ ≪ *σ*_*x*_, Eq.(C.7b) requires |*g*_*x*_| to be very small, while |*g*_*y*_| is not needed to be very small, which allows 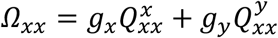 to be non-small. Therefore, analogously to the case of branching lines, distortion of a trait space affects the branching area conditions when *σ*_*y*_ ≪ *σ*_*x*_.

## Appendix D: Evolutionary branching conditions in distorted trait spaces of arbitrary higher dimensions

We derive conditions for evolutionary branching points, lines, and areas in an arbitrarily distorted trait space **s** = (*x*_1_, …, *x*_*N*_)^T^ of an arbitrary dimension *N* with *N* ≥ 2. The derivation and the obtained result are analogous to the two-dimensional case (Section 3 in the main text and Appendix C).

### D.1. Assumption for mutation

We generalize the geodesic-constant-mutation assumption for two-dimensional trait spaces (Section 3.1) as follows.

Geodesic-constant-mutation assumption (for a trait space of an arbitrary dimension):

For an arbitrary point **s**_0_ = (*x*_0,1_, …, *x*_0,*N*_)^T^ in an arbitrarily distorted trait space **s** = (*x*_1_, …, *x*_*N*_)^T^, there exist coordinates 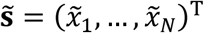 defined by

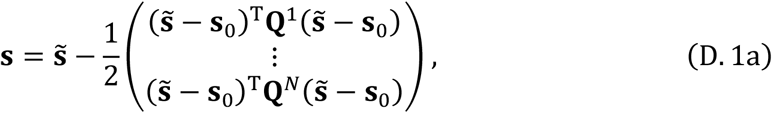

with an appropriately chosen symmetric matrices **Q**^1^,…, **Q**^*N*^, such that the mutation distribution in coordinates 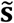 can be approximated with a constant *N*-variate Gaussian distribution,

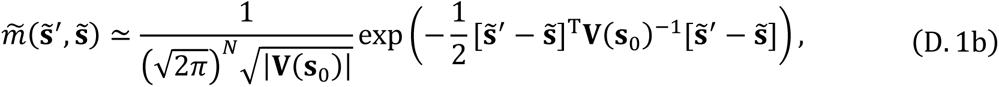

for a resident 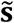 in the neighborhood of **s**_0_, satisfying

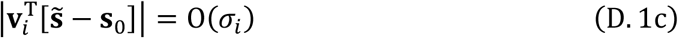

for all *i* = 1, …, *N*, with a sufficiently small *σ*_1_, …, *σ*_*N*_, where **V**(**s**_0_) has eigenvalues 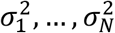 with the corresponding eigenvectors **v**_1_, …, **v**_*N*_, respectively, and *σ*_1_ ≥ ⋯ ≥ *σ*_*N*_ ≥ 0 is assumed without loss of generality.

The mutational covariance **V**(**s**) is an *N* × *N* symmetric and positive definite matrix

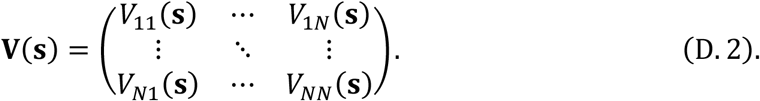

For a given **V**(**s**), we choose **Q**^*i*^ for *i* = 1, …, *N* as

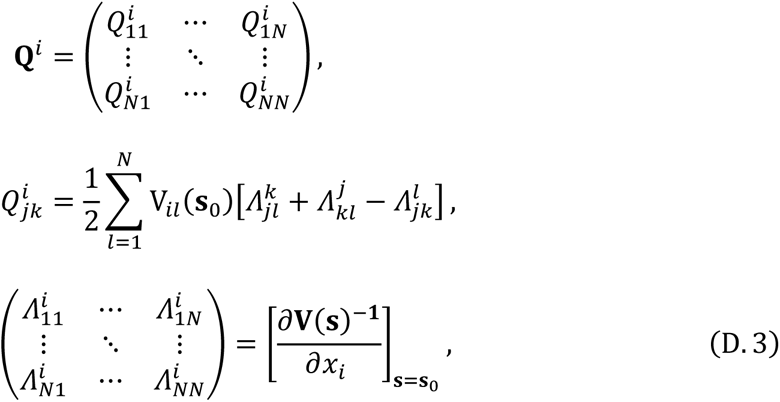

so that **V**(**s**)^−1^ has no linear dependency on 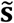 at the focal point **s**_0_ (in order to satisfy Eq. (D.1b)). In differential geometry, 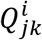 are called the Christoffel symbols of the second kind at **s**_0_ in the original coordinates **s** with respect to the metric **V**(**s**)^−1^.

### D.2. Quadratic approximation of invasion fitness functions

To reduce complexity of the expressions in the subsequent analysis, without loss of generality we assume that coordinates **s** are first rotated so that **V**(**s**_0_) become a diagonal matrix expressed as

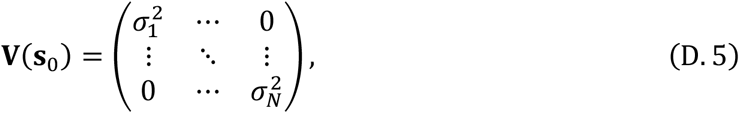

and then the geodesic coordinates 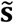 are obtained from Eqs. (D.1-3). Then, in the same manner with Eqs. (5) in the main text, we expand *f*(**s**′, **s**) around the focal point **s**_0_ as

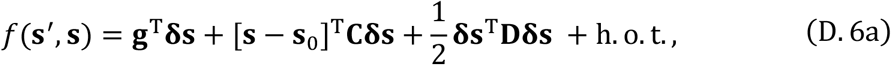

with **δs** = **s**′ − **s** and

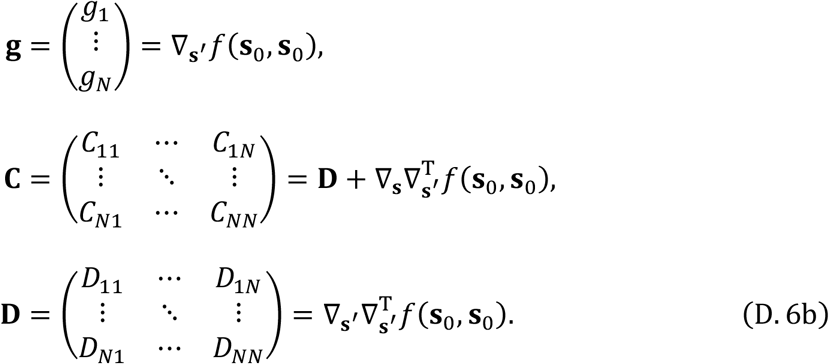

Substitution of Eq. (D.1a) into Eqs. (D.6) gives the invasion fitness function in the geodesic coordinates,

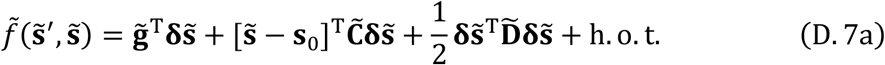

with 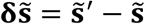 and

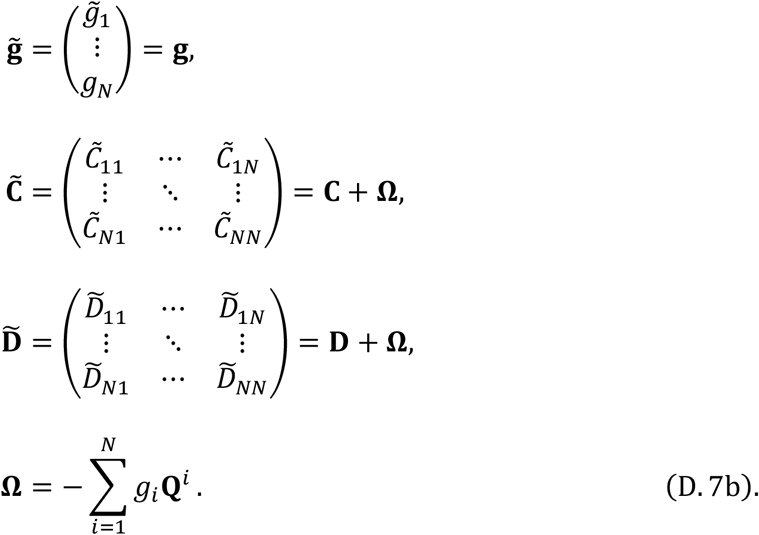

### D.3. Conditions for evolutionary branching points

In trait spaces of dimensions higher than two, it has not been formally proved yet whether points that are strongly convergence stable and evolutionarily unstable ensure high likelihoods of evolutionary branching (but see Geritz et al., 2016). Thus, such points are called candidate branching points (Ito and Sasaki, 2016). The conditions for the focal point **s**_0_ being a candidate evolutionary branching point (Ito and Dieckmann, 2014; Geritz et al., 2016; Ito and Sasaki, 2016) are described as follows.

Candidate-branching-point conditions in distorted ***N***-dimensional trait spaces:

In an arbitrarily distorted ***N***-dimensional trait space **s** = (*x*_1_, …, *x*_*N*_)^T^, a point **s**_0_ = (*x*_0,1_, …, *x*_0,*N*_)^T^ is a candidate-branching-point, if **s**_0_ satisfies the following three conditions in the corresponding geodesic coordinates 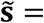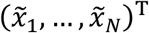 given by Eqs. (D.1–3) (after rotation of coordinates **s** so that Eq. (D.5) holds).

i. **s**_0_ is evolutionarily singular, satisfying

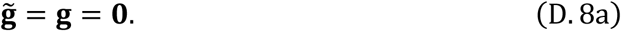
ii. **s**_0_ is strongly convergence stable, i.e., the symmetric part of

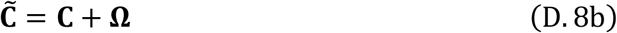

is negative definite.
iii. **s**_0_ is evolutionarily unstable, i.e., a symmetric matrix

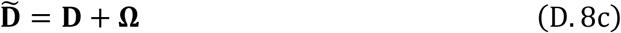

has at least one positive eigenvalue.

Since condition (i) 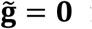 requires 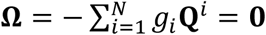, we see 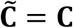 and 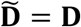. Thus, analogously to the two-dimensional case in the main text, the candidate-branching-point conditions are not affected by the distortion in trait spaces of arbitrary higher-dimensions.

### D.4. Conditions for candidate-branching-surfaces

It has not been formally proved yet whether the higher-dimensional extension of branching line conditions (Ito and Dieckmann, 2014) ensures high likelihoods of evolutionary branching. In this sense, we refer to the extended branching line conditions as the “candidate-branching-surface conditions.” If we can find an integer *L* with 1 ≤ *L* < *N* such that *σ_L_* ≫ *σ*_*L*+1_ (i.e., *σ*_*L*+1_, …, *σ_N_* are all significantly smaller than *σ*_1_, …, *σ*_*L*_), then we can simplify the original candidate-branching-surface conditions (Ito and Dieckmann 2014), in a manner analogous to the two-dimensional case (Appendix C). Consequently, we get the Candidate-branching-surface conditions for distorted trait spaces of arbitrary dimensions, described below.

Candidate-branching-surface conditions in distorted trait spaces of arbitrary dimensions (simplified):

In an arbitrarily distorted N-dimensional trait space **s** = (*x*_1_, …, *x*_*N*_)^T^, there exists an (*N* − *L*)-dimensional candidate-branching-surface containing a point **s**_0_, if **s**_0_ satisfies the following four conditions in the corresponding geodesic coordinates 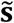 given by Eqs. (D.1–3) (after rotation of coordinates **s** so that Eq. (D.5) holds).

i. At **s**_0_ the sensitivity of 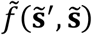 to single mutational changes of 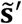 and 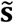 is significantly lower in a subspace 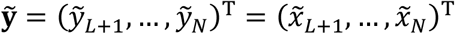 than in 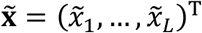, satisfying

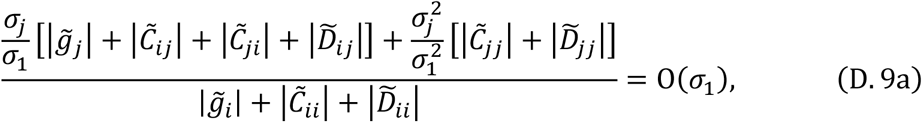

for all *i* = 1, …, *L* and *j* = *L* + 1, …, *N*, so that the normalized invasion fitness function can be simplified into

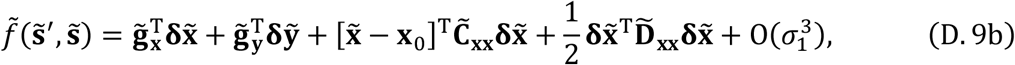

with x_0_ = (*x*_0,1_, …, x_o,l_)^T^ and

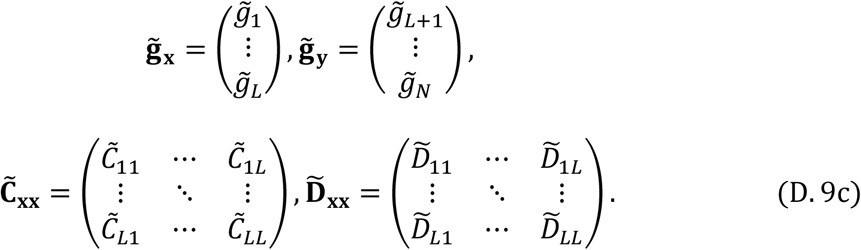
ii. **s**_0_ is evolutionarily singular in subspace 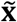, satisfying

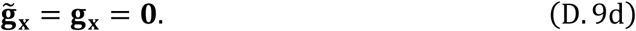
iii. **s**_0_ is strongly convergence stable in subspace 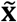, i.e., the symmetric part of

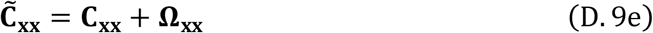

is negative definite, where

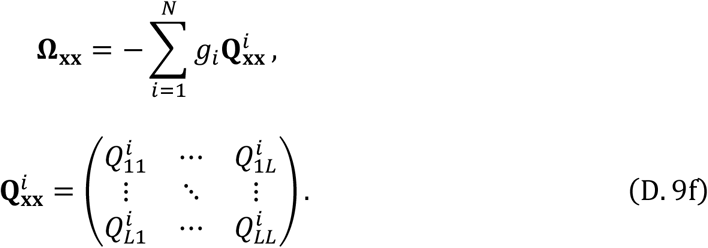
iv. **s**_0_ is sufficiently evolutionarily unstable (corresponding to disruptive selection) in subspace 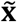, satisfying

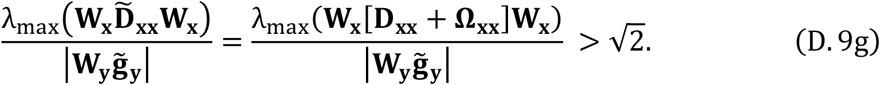

where **W**_x_ and **W**_y_ are diagonal matrices with their diagonal components *σ*_1_, …, *σ*_L_ and *σ*_L+1_, …, *σ*_N_, respectively, and λ_max_() gives the maximum eigenvalue of its argument matrix.

Note that even when condition (ii), **g**_***x***_ = (*g*_1_, …, *g*_l_)^T^ = **0**, is satisfied, ***g***_***y***_ = (*g*_*L*+1_, …, *g_N_*)^T^ can make 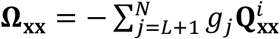 non-zero. Therefore, analogously to the two-dimensional case (Section 3.4 and Appendix C.4), the candidate-branching-surface conditions are affected by the distortion.

When the subspace 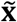 is one-dimensional (*L* = 1), the candidate-branching-surface conditions are proved to ensure evolutionary branching in the maximum likelihood invasion-event paths (Ito and Dieckmann, 2014). But for other cases (*L* > 1), those conditions only give candidates, which do not ensure high likelihoods for evolutionary branching.

When *σ_L_* ≫ *σ*_*L*+1_ is satisfied, all possible mutants are almost restricted to 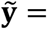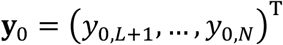, which upon substituting into Eq. (D.1a) gives an (*N* − *L*)-dimensional constraint surface expressed in coordinates **s**,

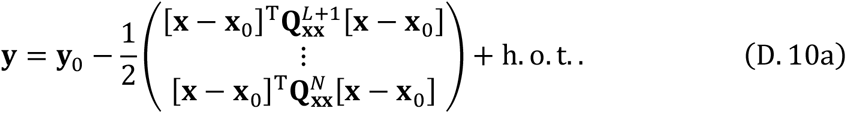

If *σ*_*L*+1_, …, *σ_N_* are all zero, then the candidate-branching-surface conditions (Eqs. (D.9)) become identical to the conditions for candidate-branching-points along a constraint surface locally described in the form of Eq. (D.10a), derived by Ito and Sasaki (2016). Specifically, we describe the constraint surface, Eq. (D.10a), as

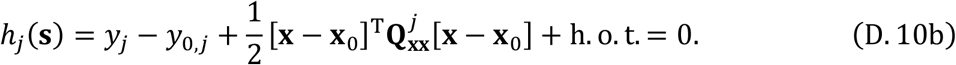

for *j* = *L* + 1, …, *N*, and define a Lagrange invasion fitness

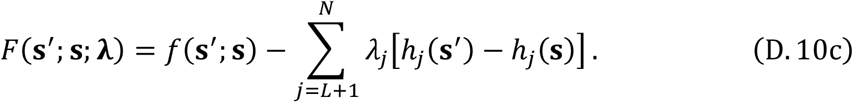

Then by Theorem 2 in Ito and Sasaki (2016), we get **λ** = (*λ*_*L*+1_, …, *λ*_*M*_)^T^ = ***g***_***y***_, and find the fitness gradient **g**_**h**_, fitness gradient variability **C**_**h**_, and fitness curvature **D**_**h**_ along the constraint surface at the focal point **s**_0_ as

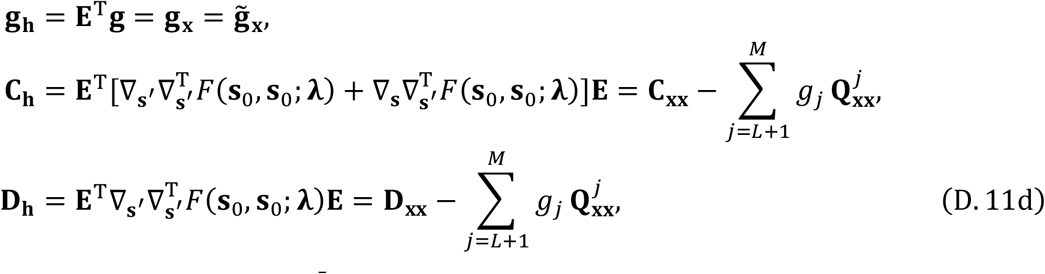

where 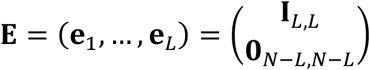 consists of the orthogonal base vectors **e**_1_, …, **e**_*L*_ of the tangent plane of the surface at **s**_0_, with I_*L,L*_ an *L* × *L* identity matrix and 0_*N-L*,*N-L*_ an (*N* − *L*) × (*N* − *L*) zero matrix. Thus, when Eq. (D9d), *g*_*x*_ = (*g*_1_, …, *g*_*L*_)^T^ = 0, holds, 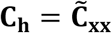and 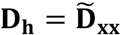 both hold. Therefore, the above candidate-branching surface conditions under *σ*_*L*+1_, …, *σ*_*N*_ = 0 are identical to the branching point conditions along (*N* − *L*)-dimensional constraint surfaces (Ito and Sasaki, 2016).

### D.5. Branching area conditions

The branching area conditions have not been developed for trait spaces of dimensions higher than two. Here we heuristically extend the simplified candidate-branching-surface conditions in Appendix D.4 into the simplified branching area conditions, in a manner analogous to the two-dimensional case. Specifically, we propose the higher-dimensional simplified branching area conditions as follows.

Branching area conditions in distorted trait spaces of arbitrary dimensions (simplified)

In an arbitrarily distorted ***N***-dimensional trait space **s** = (*x*_1_, …, *x*_*N*_)^T^, there exists an evolutionary branching area containing a point **s**_0_, if **s**_0_ satisfies the following two conditions in the corresponding geodesic coordinates 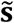 given by Eqs. (D.1–3) (after rotation of coordinates **s** so that Eq. (D.5) holds), under *σ*_1_, …, *σ_L_* ≫ *σ*_*L*+1_, …, *σ_N_*.

i. The symmetric part of

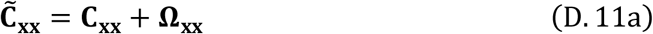

is negative definite (i.e., **s**_0_ is strongly convergence stable in subspace 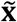 when *g*_*x*_ = 0).
ii. **s**_0_ satisfies

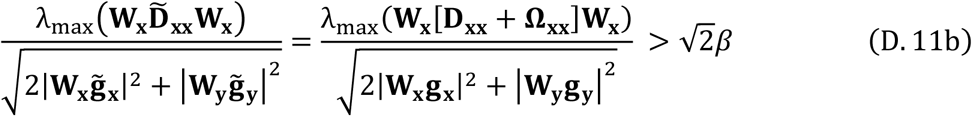

with 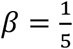 (i.e., **s**_0_ is sufficiently evolutionarily unstable in subspace 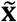 when *g*_*x*_ = 0), where **W**_x_ and **W**_y_ are diagonal matrices with its diagonal components *σ*_*L*_, …, *σ*_*L*_ and *σ*_*L*+1_, …, *σ*_*N*_, respectively, and *λ*_max_() gives the maximum eigenvalue of its argument matrix.

Under *σ*_1_, …, *σ*_*L*_ ≫ *σ*_*L*+1_, …, *σ_N_*, i.e., λ_max_(W_x_) ≫ λ_max_(W_y_), Eq.(D.11b) requires |*g*_*x*_| = |(*g*_1_, …, *g*_*L*_)^T^| to be very small, while |*g*_*y*_| = |(g_*L*+1_, …, g_*N*_)^T^| is not needed to be very small, which allows 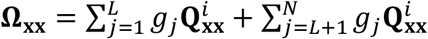 to be non-small. Therefore, analogously to the two-dimensional case in Appendix C.5, the distortion affects the branching area conditions under *σ*_1_, …, *σ*_*L*_ ≫ *σ*_*L*+1_, …, *σ_N_*, in distorted trait spaces of arbitrary dimensions.

## Appendix E: Describing directional evolution in geodesic coordinates

When the mutation distribution non-negligibly deviates from the Gaussian distribution, the canonical equation for directional evolution, Eqs. (1) in the main text, may not be warranted (Dieckmann and Law, 1996). In this case, if the geodesic-constant-mutation assumption (Section 3.1 and Appendix D.1, for two- and the higher-dimensional trait spaces, respectively) holds good, the canonical equation described in the geodesic coordinates is still warranted.

Specifically, in a trait space **s** = (*x*_1_, …, *x*_*N*_)^T^ of an arbitrary dimension N, we assume a directionally evolving population with a resident phenotype **s**. Even when the mutation distribution *m*(**s**′, **s**) cannot be approximated with an N-variate Gaussian distribution, the geodesic coordinates for the focal point **s**_0_ chosen at the resident phenotype **s** allows the approximation, if the geodesic-constant-mutation assumption holds. In this case, in the same manner with Eqs. (1), we can describe the directional evolution in the geodesic coordinates 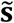 as

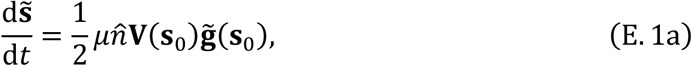

with

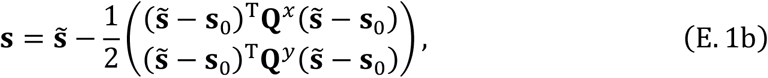

where 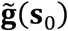 is the fitness gradient in coordinates 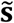. For 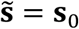, we see that

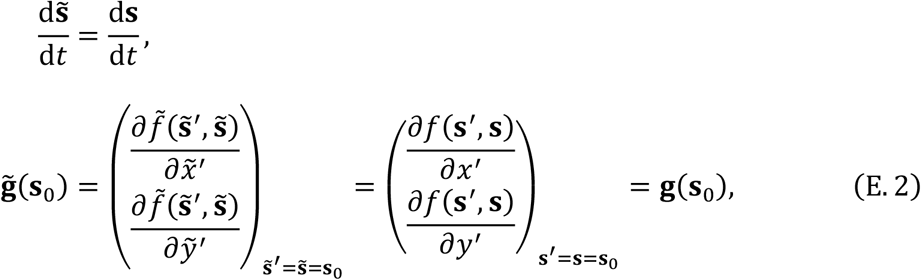

which upon substitution into Eq. (E.1a) gives

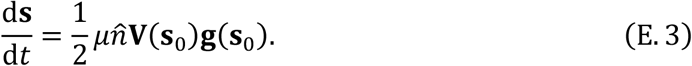

Replacing **s**_0_ with **s** gives

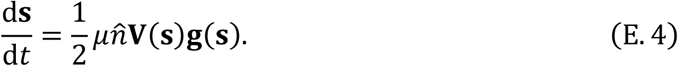

Eq. (E.4) is the same with Eq. (1a) in the main text except that **V**_*m*_(**s**) is replaced with **V**(**s**). Note that differentiation of Eq. (E.4) is not warranted. Thus, for numerical simulation by differentiation, Eq. (E.1) is better than Eq. (E.4).

## Appendix F: Analysis of evolutionary branching in Example

In the main text, the original coordinates **s** are first rotated so that its mutational covariance at the focal point becomes diagonal, and then the rotated coordinates are denoted by **s** again. To avoid confusion, only in this section we distinguish the original coordinates before the rotation and after the rotation, by calling the former the “original coordinates”, denoted by 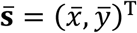, and calling the latter the “rotated original coordinates”, denoted by **s** = (*x*, *y*)^T^.

### F.1. Mutational covariance

In coordinates **u** = (*θ*, *r*)^T^, the mutational covariance is given by a constant diagonal matrix

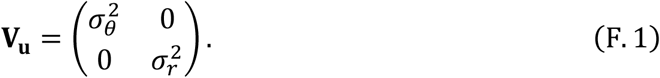

Since 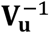 can be treated as a metric for coordinates **u**, we describe the mutational square distance from **u** to **u** + **du** with infinitesimal **du** = (d*θ*, d*r*)^T^ as

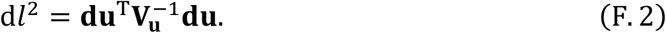

By taking the first derivative of Eqs. (26) in the main text,

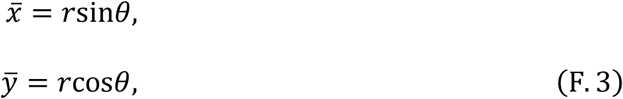

we express an infinitesimally small 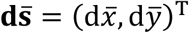 as

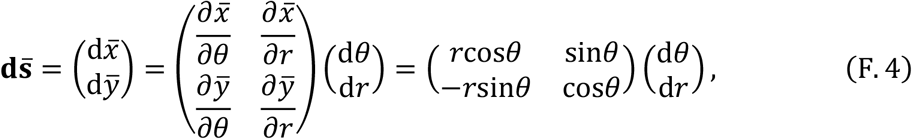

which gives

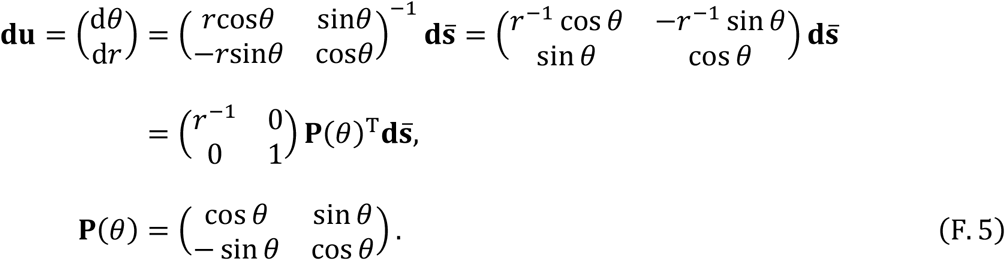

Substituting Eq. (F.5) into Eq. (F.2) gives

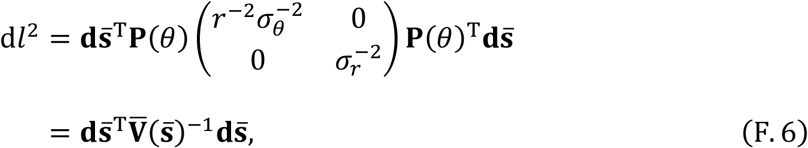

which gives the mutational metric in the original coordinates,

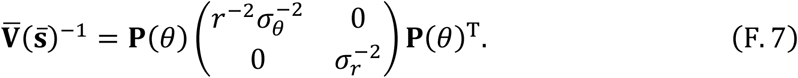

Next, we rotate the original coordinates 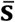 about the focal point 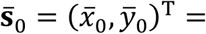 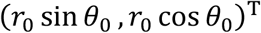 into the rotate original coordinates **s** = (*x*, *y*)^T^ by

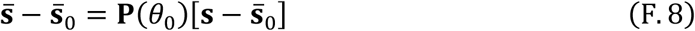

with a rotation matrix **P**(*θ*_0_). Eq. (F.8) gives 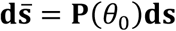, and substituting it into Eq. (F.6) gives

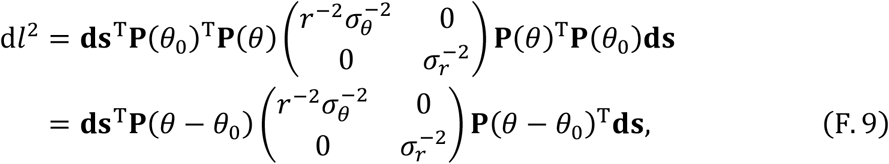

from which we get the mutational metric in the rotated original coordinates,

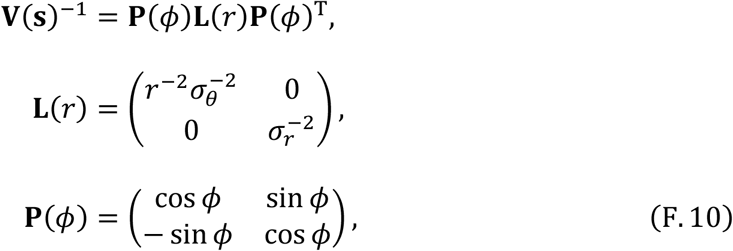

where *ϕ* = *θ* − *θ*_0_, and 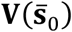 is given by

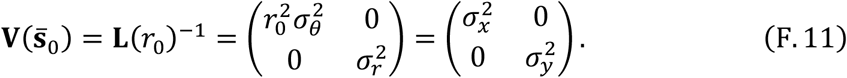

For convenience, we express 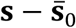 in terms of *r* and *ϕ* = *θ* − *θ*_0_, as

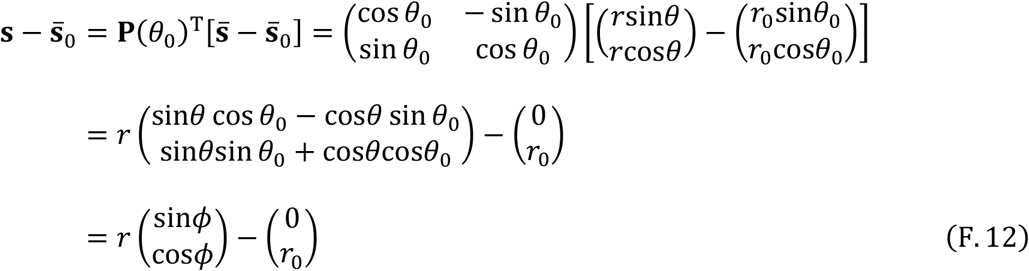

Note that the focal point 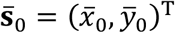 corresponds to (*ϕ*, *r*)^T^ = (*ϕ*_0_, *r*_0_)^T^ with *ϕ*_0_ = 0.

### F.2. Distortion matrices

By using Eq. (F.10), we express the first derivative of the mutational metric **V**(**s**)^−1^ as

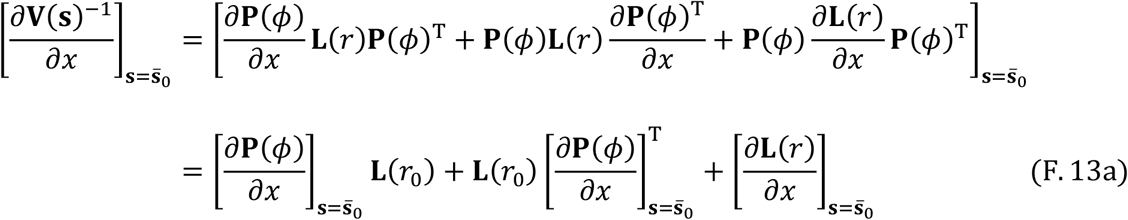

and

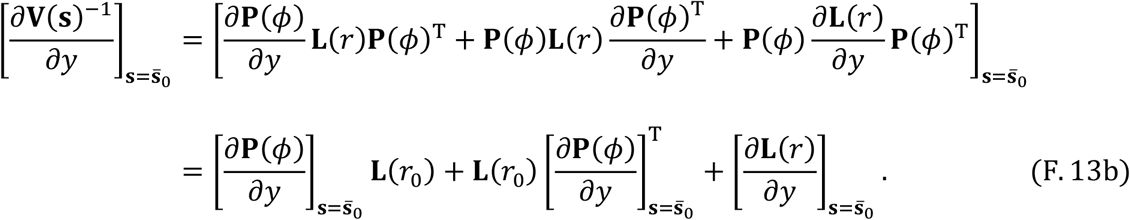

From Eqs. (F.12) we see

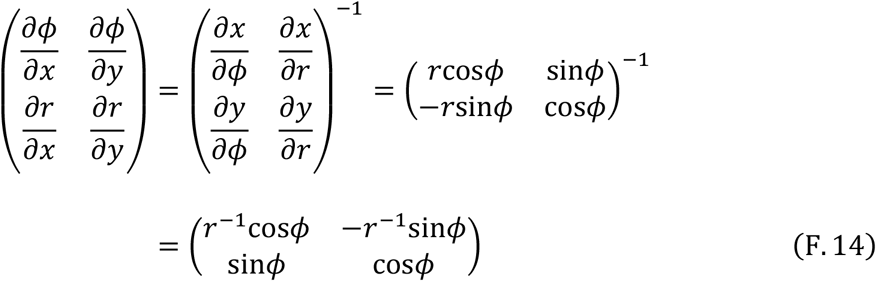

and thus

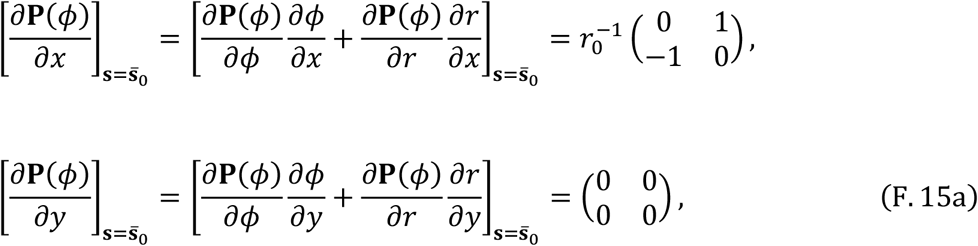

and

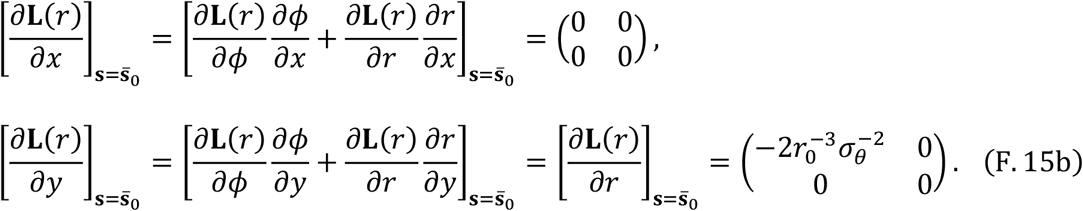

Substituting Eqs. (F.15) into Eqs. (F.13) gives

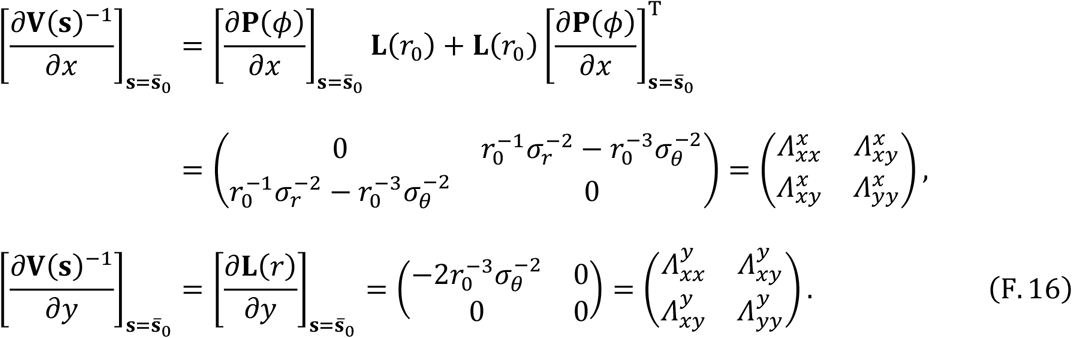

Finally, substituting Eqs. (F.11) and (F.16) into Eqs. (12) in the main text, we get

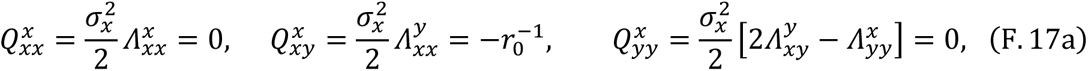

and

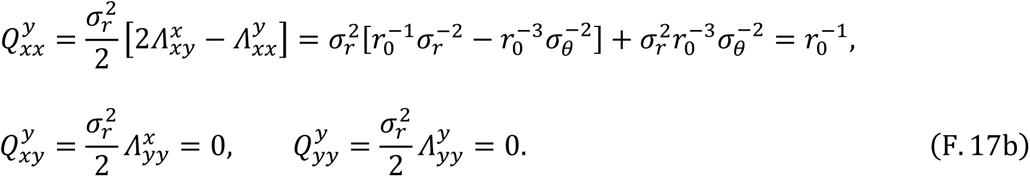

### F.3. Geodesic invasion fitness function

In the original coordinates before rotation, 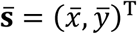, we express the invasion fitness function (Eq. (25) in the main text) as

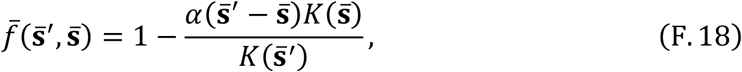

and expand it around the focal point 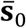 as

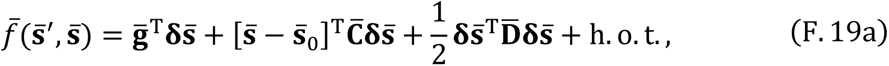

with 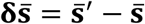 and

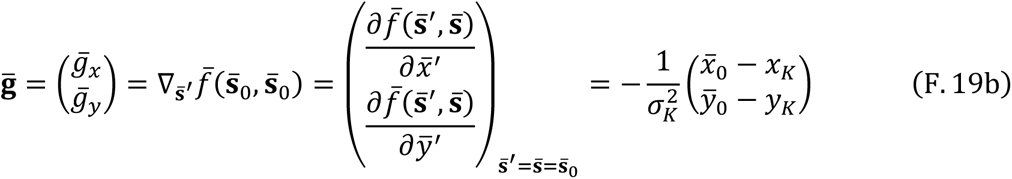

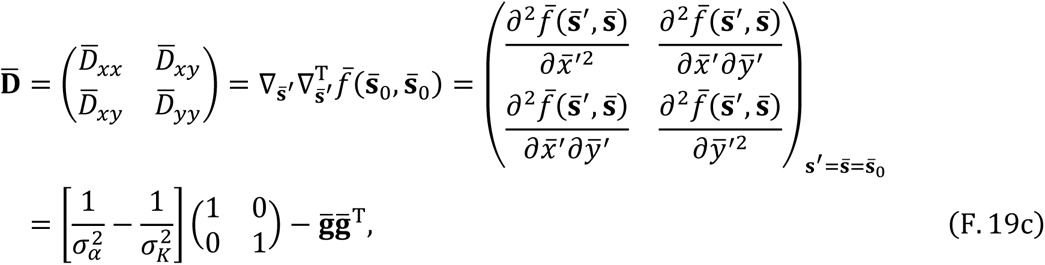

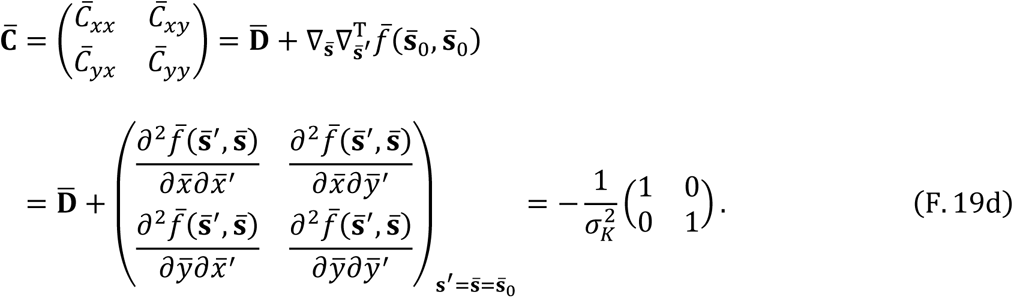

Substituting Eq. (F.8) into Eqs. (F.19) gives the invasion fitness in the rotated original coordinates **s**,

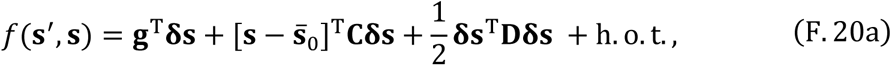

with

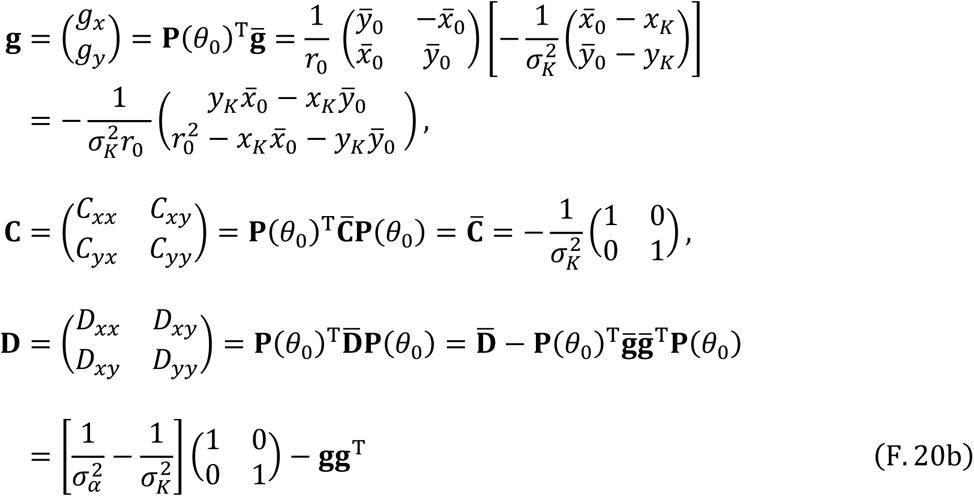

and

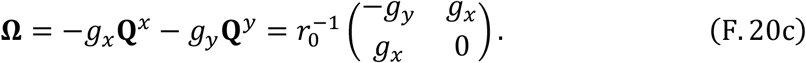

In addition, Eq. (F.11) gives

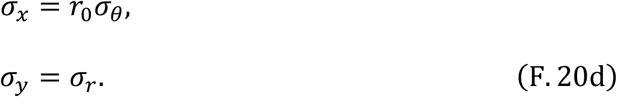

### F.4. Evolutionary branching points

Since the branching point conditions are not affected by the distortion, as shown in Section 3.3, we can directly examine those conditions in the original coordinates 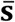 (or in the rotated original coordinates **s**, equivalently), by using Eqs. (F.19). Condition (i) (for evolutionary singularity), 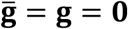, gives a unique evolutionarily singular point 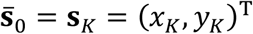. At the point, we see

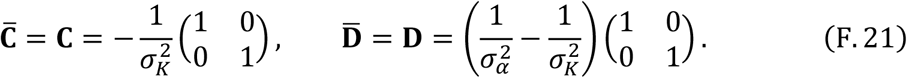

Thus, condition (ii) (for strong convergence stability) is always satisfied. Condition (iii) (for evolutionary instability) is satisfied if and only if *σ*_*α*_ < *σ*_*K*_. Therefore, a necessary and sufficient condition for existence of an evolutionary branching point is given by *σ*_*α*_ < *σ*_*K*_.

### F.5. Evolutionary branching lines

We apply the simplified branching line conditions described in Section 3.4, by substituting Eqs. (F.20) into Eqs. (21) in the main text. For simplicity, we assume that *σ*_*y*_ = *σ*_*r*_ is much smaller than *σ*_*x*_ = *r*_0_*σ*_*θ*_, so that condition (i) is satisfied. Then condition (ii) (for evolutionarily singular line) is given by

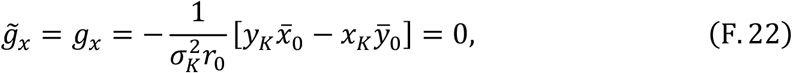

which forms a line

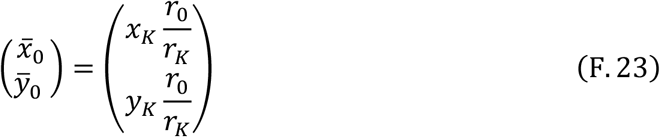

with 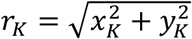 and a positive parameter *r*_0_. From Eqs. (F.20) and (F.23) we see

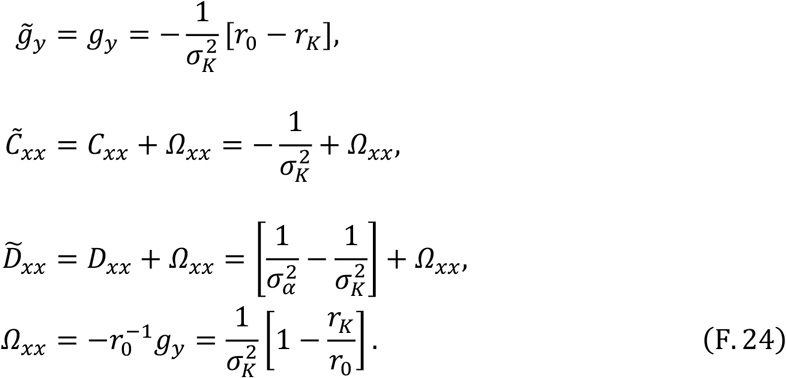

Thus, condition (ii) (for convergence stability along 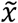) is expressed as

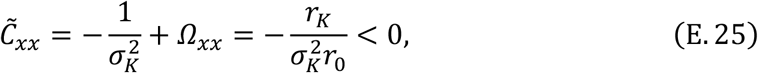

which is always satisfied because *r*_0_ > 0. Condition (iv) (for sufficient disruptive selection along 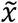) is expressed as

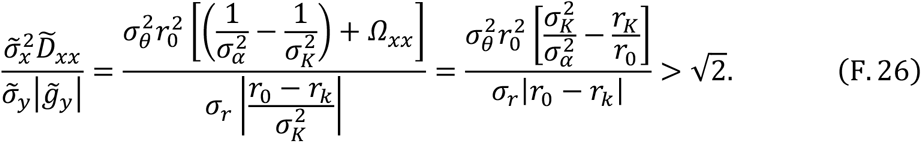

### F.6. Meaning of geodesic invasion fitness

Here we show that the geodesic invasion fitness function for a focal point describes the invasion fitness function in the undistorted coordinates **u** = (*r*, *θ*)^T^ up to the second order terms.

Substituting Eqs. (24) and (26) into Eq. (25), we express the invasion fitness in coordinates **u** = (*r*, *θ*)^T^ as

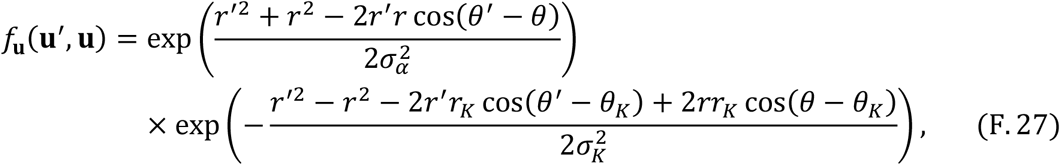

which is expanded around the point **u**_0_ = (*θ*_0_, *r*_0_)^T^ corresponding to the focal point 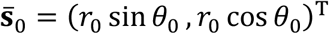 as

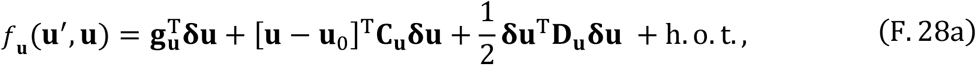

with

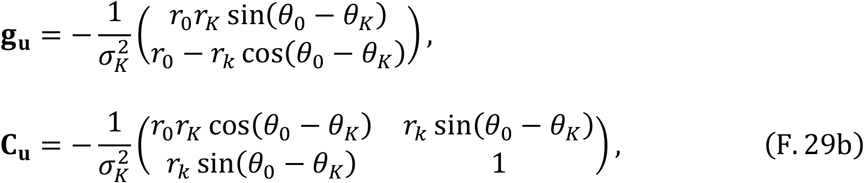

and

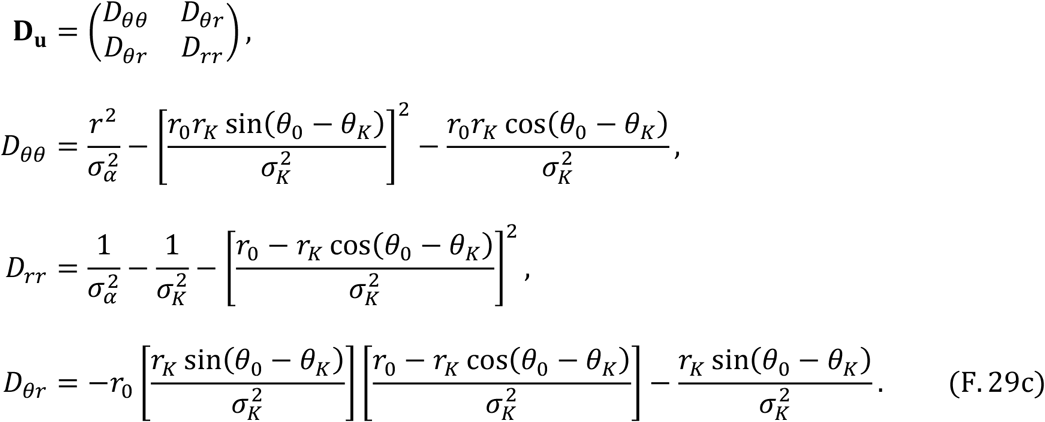

On the other hand, we can express the geodesic invasion fitness (from Eqs. (18) with Eqs. (F.20)) as

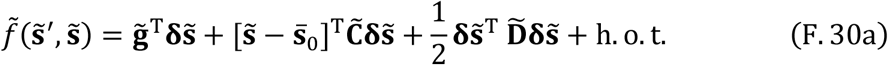

with

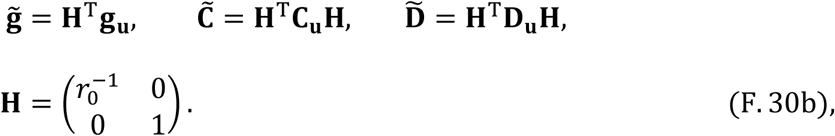

Note that the coordinates **u** = (*r*, *θ*)^T^ has a globally constant mutational covariance 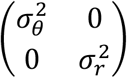, while the geodesic coordinates 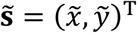 have a locally constant mutational covariance 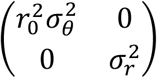 around the focal point 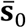. Thus, we scale 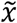 of the geodesic coordinates by 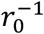 (and shift 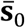 to **u**_0_), by introducing new coordinates

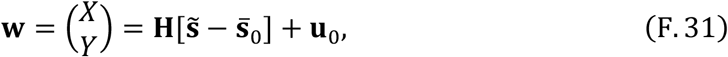

to attain the same covariance matrix 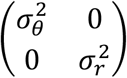 with that of the coordinates **u** = (*r*, *θ*)^T^. Then we get the scaled geodesic invasion fitness,

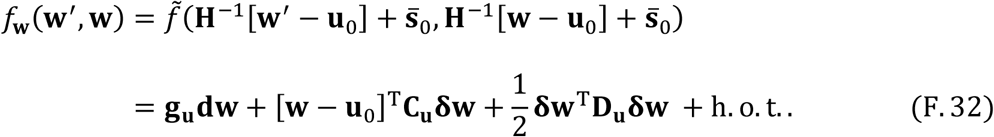

Note that Eq. (F.32) is identical to Eq. (28a). Therefore, the scaled geodesic fitness function *f*_**w**_(**w**′, **w**) describes the invasion fitness function *f*_**u**_(**u**′, **u**) in the non-distorted coordinates **u** = (*r*, *θ*)^T^ up to the second order terms. Since all of the conditions for evolutionary branching points, lines, and areas in this paper concern only the first and second order derivatives of invasion fitness functions, application of those conditions in the coordinates **u** = (*r*, *θ*)^T^ give identical results to those in the scaled geodesic coordinates **w** = (*X*, *Y*)^T^, which are also identical to those in the geodesic coordinates 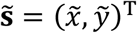, because a linear coordinate transformation does not affect those conditions.

## Appendix G: Simulation algorithm for evolutionary dynamics

This study conducts numerical simulation of evolutionary dynamics in Example as trait substitution sequences based on the oligomorphic stochastic model defined by Ito and Dieckmann (2014). The oligomorphic stochastic model in Ito and Dieckmann (2014) is the same with the algorithm described in Ito and Dieckmann (2007), except that population dynamics after each mutant invasion is directly calculated in Ito and Dieckmann (2014). The algorithm of the oligomorphic stochastic model used in this study is listed below.

0. [Initial setting] Set initial M phenotypes **s**_1_, …, **s**_*M*_ at time *t* = 0 (This study uses *M* = 1, corresponding to an initially monomorphic community). Calculate equilibrium population densities 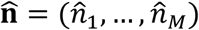 at which d*n*_*k*_ / d*t* = 0 for all *k* = 1, …, *M*. Define the extinction threshold *ε*.
1. [Mutant emergence] Choose resident *i* with probability *w_i_/w*, where 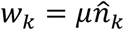 is the emergence rate of a mutant from resident **s**_*K*_, with *μ* the mutation rate per unit population density per one generation, and 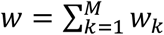. Choose a mutant 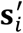 according to the mutation distribution 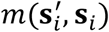.
2. [Time updating] Update time *t* by adding 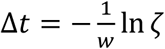, where 0 < *ζ* ≤ 1 is a uniformly distributed random number.
3. [Mutant invasion] Choose a uniformly distributed random number 0 < *ζ* ≤ 1. If *ζ* is smaller than the invasion fitness 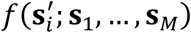 of the mutant phenotype 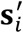 against residents **s**_1_, …, **s**_*M*_ at 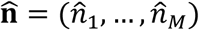, proceed to Step 4. Otherwise, return to Step 1.
4. [Population dynamics triggered by mutant invasion] Increase *M* by 1 and set **s**_*M*_ = **s**′. Calculate equilibrium population densities from population dynamics with initial population densities 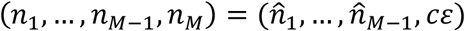 with a constant c ≥ 1. In the course of these population dynamics, delete phenotypes **s**_*K*_ with *n_k_* < *ε*, and decrease *M* accordingly.
5. Continue with Step 1.

Note that the time taking for population dynamics triggered by a mutant invasion to reach the next equilibrium (Step 4) is assumed to be negligible in comparison with waiting times for invading mutants (Step 2). The above algorithm is slightly simplified from Ito and Dieckmann (2014), by assuming that the birth rate per unit population density per unite time is equal to 1.

For the numerical simulation in Example, the two parameters *ε* and *c* are set at *ε* = 1 × 10^−1^ and *c* = 10^3^. Occurrence of evolutionary branching are treated as emergence of polymorphic residents with the maximum distance among them exceeding 15 × *σ*_*θ*_ along *θ*.

